# Direct pathway bias and altered striatal neurogenesis in human iPSC models of 16p11.2 CNVs: Evidence from single-cell and functional analyses

**DOI:** 10.1101/2025.02.26.640425

**Authors:** Marija Fjodorova, Parinda Prapaiwongs, Olena Petter, Ilaria Morella, Zongze Li, Jessica Hall, Ngoc-Nga Vinh, Sara Davies, Jonathan Green, Jeremy Hall, Marianne van den Bree, Riccardo Brambilla, Meng Li

## Abstract

Striatal medium spiny neurons (MSNs) control motor, cognitive, and social domains via direct (dMSNs) and indirect (iMSNs) basal ganglia pathways. Recent genomic analyses implicate striatal circuit dysfunction in neurodevelopmental disorders (NDDs) and highlight MSNs as a newly recognised cell type affected in schizophrenia, yet much NDD research still focuses on cortical interneurons and glutamatergic neurons, leaving MSN involvement understudied. Here, we use human iPSC-derived MSNs to demonstrate high-fidelity striatal development and explore 16p11.2 copy number variants (CNVs) – mutations predisposing carriers to autism spectrum disorder, schizophrenia, intellectual disability and other NDD conditions featuring basal ganglia deficits. By profiling both 16p11.2 duplication and deletion MSNs, we uncover reciprocal changes in MSN neurogenesis kinetics that converge on a dMSN fate bias. These shifts correspond to altered calcium signalling and enhanced firing upon direct-pathway activation. Our findings reveal a previously unappreciated role for MSN subtype imbalances in NDD pathogenesis and open new avenues for therapeutic intervention.

## Introduction

During striatal development, medium spiny neurons (MSNs) diversify into two major subtypes – direct pathway (dMSNs) and indirect pathway (iMSNs) – which form the basis of basal ganglia circuitry essential for cognitive, motor, and social behaviours. The early dysfunction and subsequent loss of MSNs is a hallmark of Huntington’s disease^1^. However, a growing body of genomic studies suggests a role for basal ganglia circuits in psychiatric and neurodevelopmental disorders (NDDs), including schizophrenia, autism spectrum disorder (ASD), intellectual disability, bipolar disorder, and depression^2–5^. Moreover, in addition to glutamatergic projection neurons and cortical interneurons, MSNs were identified as a new neuronal cell type affected by schizophrenia risk variants in gene enrichment studies^6,7^. This finding may underlie certain NDD-associated cognitive and motor behavioural alterations reliant on the integrity of dMSNs^8,9^. Nonetheless, a direct causal link between human MSN deficits and NDD pathophysiology has yet to be established.

Genetic mutations disrupting basal ganglia function increase the risk of NDDs, exemplified by copy number variants (CNVs) in the 16p11.2 region (BP2–BP3 and BP4–BP5 intervals). Both deletions (16p11.2del) and duplications (16p11.2dup) are linked with NDDs, including ASD, intellectual disability and attention deficit hyperactivity disorder (ADHD), with duplications also associated with schizophrenia and bipolar features^10–13^. Rodent models of these CNVs frequently present with behavioural abnormalities observed in human carriers, such as hyperactivity, repetitive behaviours, deficits in movement control and reward learning^14–18^, consistent with cortico-striatal dysfunction. However, most NDD research in human 16p11.2 and other CNV models still focuses on GABAergic interneurons and cortical neurons^19–29^, leaving MSN involvement understudied.

Generating high fidelity human MSNs from pluripotent stem cells (PSCs) is crucial for modelling striatal development and elucidating NDD pathophysiology. Two principal approaches exist: (i) direct patterning with Activin A, a TGFβ-family ligand that specifically induces MSN progenitor fate^30–32^; or (ii) Sonic hedgehog (SHH) activation, which yields broader subpallial fates^33–36^. In addition to producing MSNs from the lateral ganglionic eminence (LGE), SHH-based methods inadvertently generate GABAergic interneurons and cholinergic neurons from the adjacent MGE, thus reducing MSN yield. By contrast, Activin A-driven protocols bypass SHH activation and restrict MGE gene expression^31^. However, previous assessments relied mainly on immunostaining or bulk transcriptome profiling^31,32,37,38^, leaving the full extent of MSN subtype recapitulation uncharacterised.

In this study, we employ single-cell RNA sequencing (scRNA-seq) to map the transcriptional landscape of Activin A-patterned MSNs against human foetal and adult striatal signatures^39–41^, confirming a high-fidelity match to *in vivo* cell populations. We then examine the impact of 16p11.2 duplications or deletions on MSN development and function in induced PSCs (iPSCs) derived from CNV carriers. Our results reveal reciprocal changes in MSN differentiation kinetics between the two 16p11.2 genotypes, but a convergent bias on one MSN subtype that translates into distinctive calcium signalling and hyperexcitability in MSNs of these carriers. This work provides new mechanistic insights into how 16p11.2 mutations may disrupt striatal circuits, contributing to motor and cognitive symptoms in CNV carriers and rodent models^15,16,42^. It further underscores the suitability of iPSC-based striatal models for studying NDD pathogenesis and identifying potential therapeutic targets.

## Results

### • Activin A patterning produces high-purity LGE identity and recapitulates human striatal signatures

We employed a droplet-based scRNA-seq platform to profile iPSC-derived striatal neurons from 13 days *in vitro* (DIV) to 40DIV. This timeframe follows anterior neuroectoderm induction, striatal fate specification, and peak MSN production in the Activin A model (**Fig 1A**)^31,32^. After quality control and batch effect correction with harmony (**Fig S1A and S1B**), we obtained transcriptomes of 59,615 single cells (**Fig 1B and S1C**). Gene expression-based prediction of cell cycle state reflected MSN differentiation kinetics where proliferating progenitors in S and G2M phases dominated at 13DIV, while non-dividing cells in G1 phase became prevalent at 18– 40DIV (**Fig S1D and S1E**).

**Figure 1.**
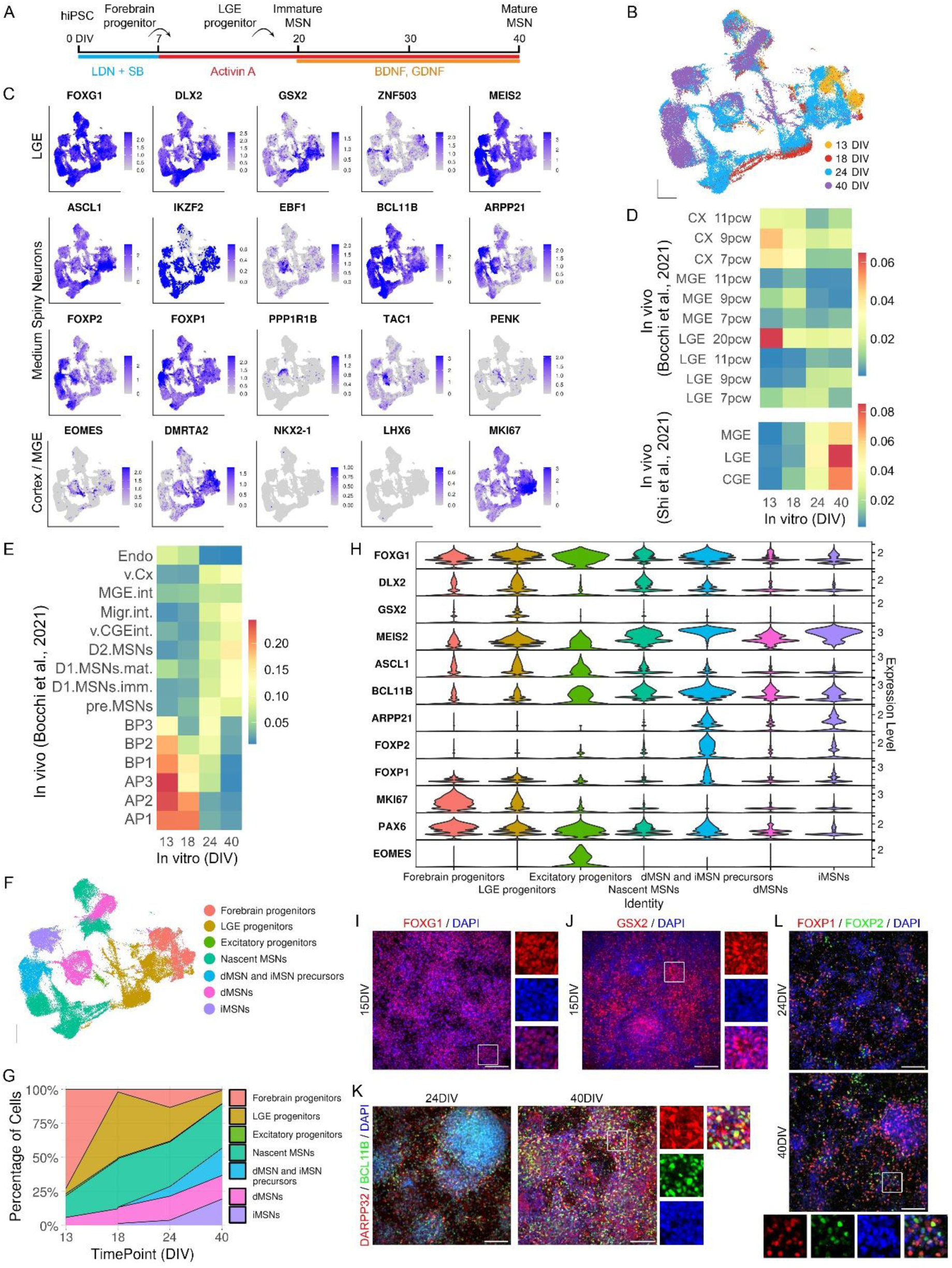
Activin A protocol yields high purity LGE identity in hiPSCs. (A) Schematic of MSN differentiation protocol. (B) UMAP embedding of scRNA-seq data of 59,615 cells from iPSC-derived MSNs at 13–40DIV. (C) Feature plots showing the expression levels of brain-region-specific genes in MSNs. (D and E) Heatmaps of the (D) region and (E) cell-type similarity between iPSC-derived MSNs and the human foetal brain (7–20PCW)^39,40^. (F) Annotation of cell types one the basis of gene expression visualised by UMAP. (G) Stack plot depicting quantification of cell population ratios at each time point. (H) Violin plot showing the expression of cell-type-specific genes in each cluster group. (I–L) Immunostaining of MSNs for FOXG1 (I), GSX2 (J), DARPP32 and BCL11B (K), FOXP1 and FOXP2 (L). Scale bars: 100 μm in (I–L). See also **Figure S1** and **Table S1**.

Regional identity of the cells was confirmed by strong expression of LGE genes including those that mark striatal progenitors (*DLX2*, *GSX2*) and MSNs (*ASCL1*, *FOXP1*, *FOXP2*, *BCL11B* and *DARPP32*) (**Fig 1C**). Minimal expression of cortical and MGE markers verified accurate striatal fate specification (**Fig 1C**). Using Jaccard similarity index method to compare our data with published bulk RNA-seq datasets from foetal forebrain regions^39,40^, we confirmed that most iPSC-derived cells mapped to LGE-specific transcriptional programmes (**Fig 1D**). Further comparisons with foetal LGE single-cell profiles showed a high degree of overlap in gene signatures^39^, with earlier time points correlating with progenitor cluster identities and later time points corresponding to maturing MSN populations (**Fig 1E and S1F**).

To classify iPSC-derived cell types, we analysed cluster-specific gene enrichment for cell type markers from previously annotated reference atlases of various primary human brain regions (**Fig 1F, S1G–S1I, Table S1**)^39,40^. Results showed that most cells were striatal although a small cluster of cortical excitatory progenitors was observed (**Fig 1F–1H**), consistent with our previous report^31^. Cells at earlier time points largely belonged to forebrain/LGE progenitor groups, whereas nascent MSNs and dMSN/iMSN precursors expressing *BCL11B*, *MEIS2*, *FOXP1/2* emerged by 18– 24DIV, and mature MSN subtypes were detectable at 24–40DIV (**Fig 1G and 1H**). These transcriptomic findings were corroborated by abundant FOXG1^+^ and GSX2^+^ progenitors from 15DIV onward and BCL11B^+^, DARPP32^+^, FOXP1^+^, FOXP2^+^ MSNs from 24DIV (**Fig 1I–L**). We also found that striatal fate and MSN subtype neurogenesis were induced more rapidly in Activin A-treated iPSCs than SHH-patterned cultures (**Fig 1G, S1J and S1K**)^43^.

We next assessed the authenticity and maturity of iPSC-derived MSNs by comparing our dataset with three striatal single-cell atlases encompassing foetal (7–11 and 9– 18 post-conception week [PCW]) and adult (32–83 years) human brain samples (**Fig 2A**)^39–41^. Projecting these references onto our data confirmed that iPSC-derived cells spanned the continuum from forebrain/LGE progenitors to nascent and mature dMSN/iMSN subtypes (**Fig 2B, 2C, S2A and S2B**), with foetal samples from later ages mapping to more mature MSN clusters (**Fig 2D and 2E**). In contrast, adult striatal cells preferentially aligned with the most mature iPSC-derived MSN clusters (**Fig 2B, 2C, 2F and S2C**). Reversing the projection (*in vitro* cells onto Bocchi *et al.*‘s dataset) confirmed that iPSC-derived striatal cell types corresponded closely to those *in vivo*, with MSN clusters maturing in parallel to older foetal time points (**Fig S2D–S2F**). Pseudotime indicated that MSNs developed from forebrain/LGE progenitors and branched into dMSNs/iMSNs at ∼18–24DIV, with production of dMSNs preceding birth of iMSNs as observed *in vivo* (**Fig 2G and 2H**).

**Figure 2.**
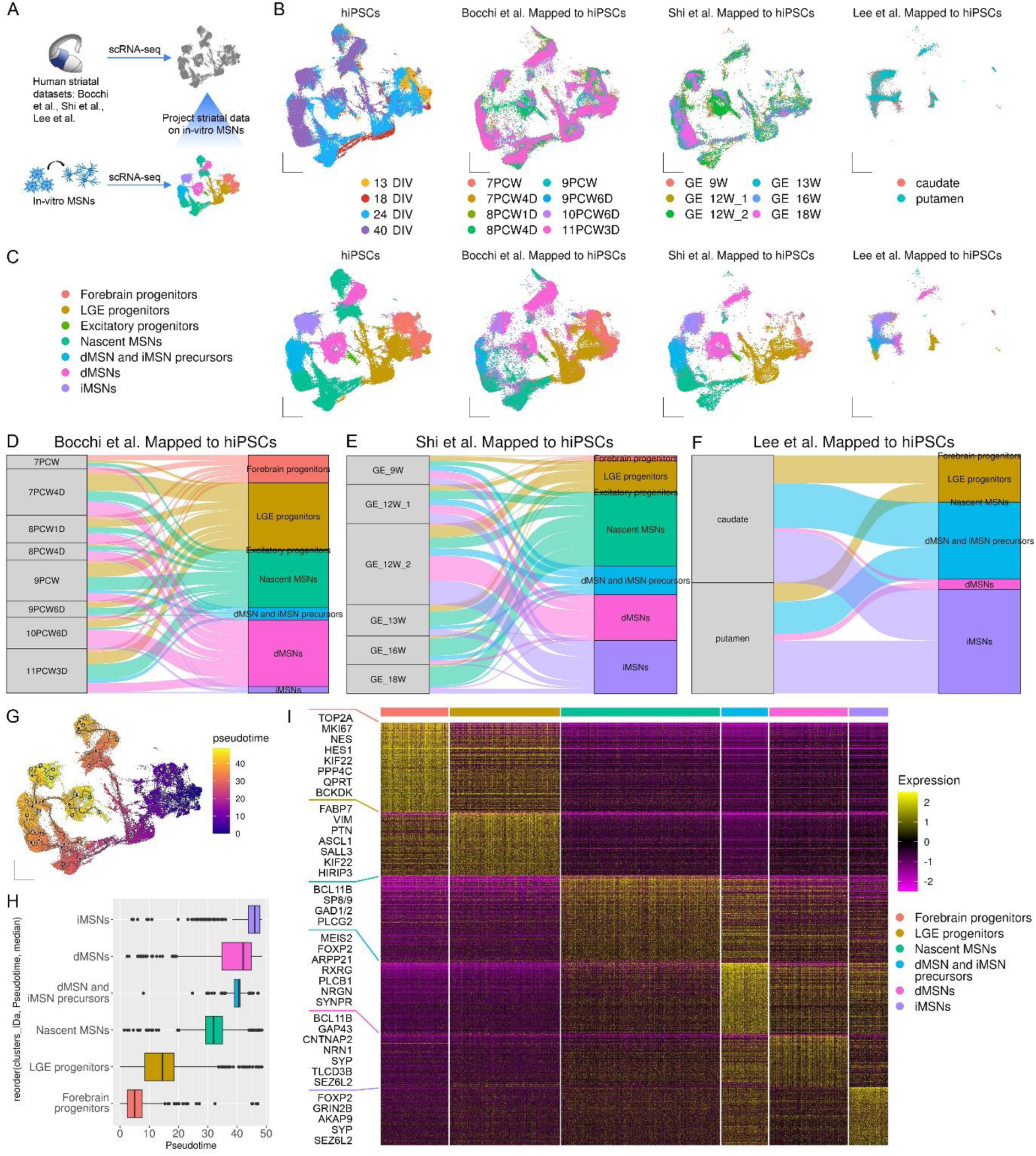
Activin A-patterned MSNs recapitulate human striatal signatures with high fidelity. (A) Overview of how human foetal^39,40^ and adult^41^ striatal cells were used to compare similarity with iPSC-derived MSNs. (B and C) UMAP embedding of MapQuery projections of striatal single-cell atlases onto scRNA-seq data coloured by (B) sample and (C) cell type. (D–F) Alluvial plots showing alignment of human striatal cells at (D) 7–11 PCW^39^, (E) 9–18 PCW^40^ or (F) 32–88 years^41^ with cell types in iPSC-derived MSNs. (G) Trajectory analysis showing the pseudotime of striatal cells on UMAP. (H) Box plot showing the pseudotime of cell types in iPSC-derived MSNs. (I) Heatmap showing DEGs enriched in each MSN cluster group. See also **Figure S2** and **Table S2**.

Finally, an integrated transcriptomic analysis of iPSC-derived MSNs and foetal striatal neurons identified conserved markers across LGE progenitors (*GSX2*, *ASCL1*) and nascent MSNs (*MEIS2*, *BCL11B*, *FOXP1/2*) (**Fig S2G–S2I**). Importantly, known dMSN genes (*EBF1*, *TAC1*, *ZFHX3*, *GAP43*) and iMSN markers (*SIX3*, *ZFHX4*) were similarly conserved across *in vitro* and *in vivo*^39,40,44^. We next detected genes that best distinguished each iPSC-derived striatal cell population by performing differential gene expression analysis. Results confirmed known LGE progenitor markers (*FABP7*, *VIM*, *SALL3*, *ASCL1*), nascent MSN genes (*BCL11B*, *SP8/9*, *GAD1/2*), and mature subtype markers (*ARPP21*, *SIX3*, *ZFHX3/4*, *GAP43*), as well as identified *NRGN* (Neurogranin) and *SYNPR* (Synaptoporin) as candidate new markers for MSN subtype precursors, and *NRN1* (Neuritin 1) for dMSNs (**Fig 2I, Table S2**). These findings demonstrate that our iPSC-derived MSNs accurately recapitulate molecular hallmarks of foetal and adult striatal neurons.

### • Altered developmental kinetics during MSN differentiation of iPSCs carrying 16p11.2 CNVs

Given the implicated role of basal ganglia in 16p11.2-associated NDDs and MSNs being identified *in silico* as a new neuronal type targeted by schizophrenia risk mutations, we examined the effect of these CNVs on MSN differentiation. Specifically, we used iPSC lines derived from three control subjects and three carriers each with 16p11.2dup or 16p11.2del (**Fig S3A and S3B**).

We observed striking opposing changes in CNV MSN differentiation kinetics, with consistently increased proportions of GSX2^+^ LGE progenitors in 16p11.2del cultures from early time points (15DIV) and reduced numbers of this population in 16p11.2dup cultures compared to controls (**Fig 3A and 3B**). Notably, FOXG1^+^ forebrain progenitors remained unaffected at 15–20DIV, confirming normal neural induction (**Fig S3C and S3D**). Reduced LGE pools in 16p11.2dup samples were accompanied by concomitant increase in cells labelled with MSN-defining markers BCL11B and DARPP32, while 16p11.2del cultures consistently displayed fewer of these cells compared to controls (**Fig 3C–3E**). Similarly, reciprocal changes emerged in ASCL1^+^ nascent MSNs at 20–25DIV (**Fig S3E and S3F**), and in FOXP1^+^ (but less so FOXP2^+^) MSNs between 15DIV and 40DIV (**Fig S3G–I**).

**Figure 3.**
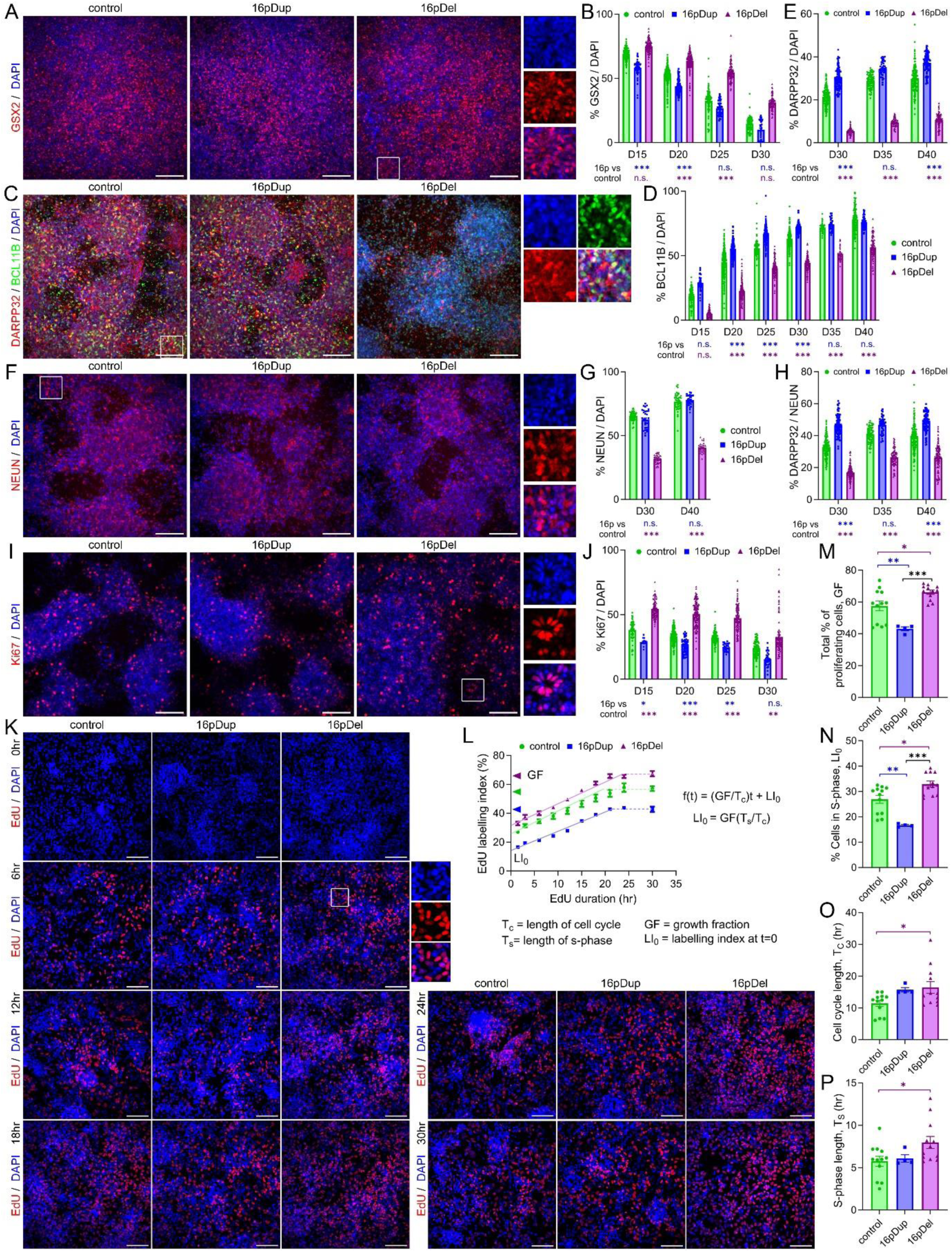
Divergent striatal neurogenesis phenotypes in MSNs carrying 16p11.2 CNVs. (A) Immunostaining and (B) quantification of differentiating control and 16p11.2dup/del iPSCs for GSX2^+^ cells. (C) Immunostaining of MSNs for BCL11B and DARPP32 with quantification of (D) BCL11B^+^ and (E) DARPP32^+^ cells. (F) Immunostaining and (G) quantification of NEUN^+^ cells. (H) Quantification of DARPP32^+^/NEUN^+^ cells in cultured MSNs. (I) Immunostaining and (J) quantification of Ki67^+^ cells in striatal cultures. (K) EdU staining and (L) quantification of EdU^+^ cells after cumulative EdU labelling for 30hr at 16DIV. (M–P) Bar plots showing cell cycle parameters calculated using the Nowakowski equation^45^ for control and 16p11.2dup/del striatal progenitors including (M) the growth fraction (GF) as indicated by dashed lines and colour coded arrowheads in (L); (N) the percentage of cells in S-phase (LI_0_) as shown in (L); (O) cell cycle length, T_C_; (P) length of S-phase, T_S_. (B,D, E, G,H, J,L and M–P) Data represent the mean ± SEM with sample sizes summarised in **Table S4**. (B,D,E,G,H and J) Bonferroni corrected multiple comparison analysis with Kruskal-Wallis test and (M–P) one-way ANOVA followed by Tukey HSD correction were used for statistical analyses. *p < 0.05, **p < 0.01, ***p < 0.001. Scale bars: 100 μm in (A,C,F,I and K). See also **Figure S3** and **Table S4**.

We also observed a reduction in NEUN^+^ cells in 16p11.2del cultures compared to controls at 30–40DIV (**Fig 3F and 3G**). Despite the global reduction of neurons in 16p11.2del cultures, significant differences in DARPP32^+^/NEUN^+^ cells persisted in both CNVs relative to controls (**Fig 3H**). Together, these data demonstrate 16p11.2-dependent preferential changes in MSN generation, albeit in opposite directions.

### • Deficits in cell cycle kinetics and proliferation underlie 16p11.2-associated striatal neurogenesis phenotypes

To investigate whether the observed changes in MSN production may be due to distorted MSN neurogenesis caused by cell-cycle perturbations, we quantified Ki67^+^ proliferating progenitors by immunocytochemistry. We observed increased Ki67^+^ cells in 16p11.2del and fewer Ki67^+^ cells in 16p11.2dup cultures from 15DIV to 30DIV (**Fig 3I and 3J**). We next measured cell-cycle parameters via cumulative 5-ethynyl-2-deoxyuridine (EdU) labelling at 16DIV (**Fig 3K and 3L**) – when GSX2^+^ and Ki67^+^ cells were produced at peak (**Fig 3B and 3J**)^45^.

In line with expanding proliferative progenitors, 16p11.2del cultures exhibited increases in both the total percentage of proliferating cells, calculated as growth fraction (GF: ∼66% vs. 58% in controls), and the percentage of cells in S-phase, as indicated by the intercept of EdU labelling curve with the y axis (LI_0_: ∼33% vs. 27%) (**Fig 3L–3N**). The total cell cycle (*T_C_*) and S-phase (*T_S_*) were also longer in 16p11.2del cells compared to controls (16.5hr and 8.0hr vs. 11.4hr and 5.8hr) (**Fig 3O and 3P**). Given that changes in G1- or S-phase can influence neuron production ^46^, this finding is consistent with a hyperproliferative state that delays neurogenic commitment. Conversely, 16p11.2dup progenitors displayed a reduction in both cycling fractions (GF: ∼43%; LI_0_: ∼17%) and a mildly increased *T_C_* (15.7hr) without changes in *T_S_* (6.1hr) (**Fig 3L–3P**), suggesting accelerated neuron production ^46^. These data explain how opposite changes in cell-cycle dynamics yield divergent neurogenesis timelines in 16p11.2 MSNs.

### • Bias toward dMSN fate in 16p11.2 cultures

To further dissect 16p11.2-dependent striatal phenotypes, we performed scRNAseq on all six 16p11.2 CNV lines and two control lines at 24DIV and 40DIV of MSN differentiation (**Fig 4A, 4B and S4A–C**). Unsupervised clustering of the 109,200 qualifying single cells revealed 22 cell populations, grouped into LGE progenitors, nascent MSNs, dMSN/iMSN precursors, dMSNs, iMSNs, plus a small excitatory neuron group (**Fig 4C, Fig S4D–S4F and Table S3**). Confirming results from EdU and Ki67 labelling assays, cells at 24DIV predicted as in S or G2/M cell cycle phases or annotated as LGE progenitors were increased in 16p11.2del samples, while the opposite direction of change was observed in 16p11.2dup cultures compared to controls (**Fig 4D, S4G and S4H**). Moreover, our analysis revealed an unexpected bias for dMSN production at 40DIV, which intriguingly occurred in both 16p11.2 genotypes (48%/5% dMSN/iMSN in duplication; 54%/3% in deletion vs. 26%/13% in controls) (**Fig 4D**).

**Figure 4.**
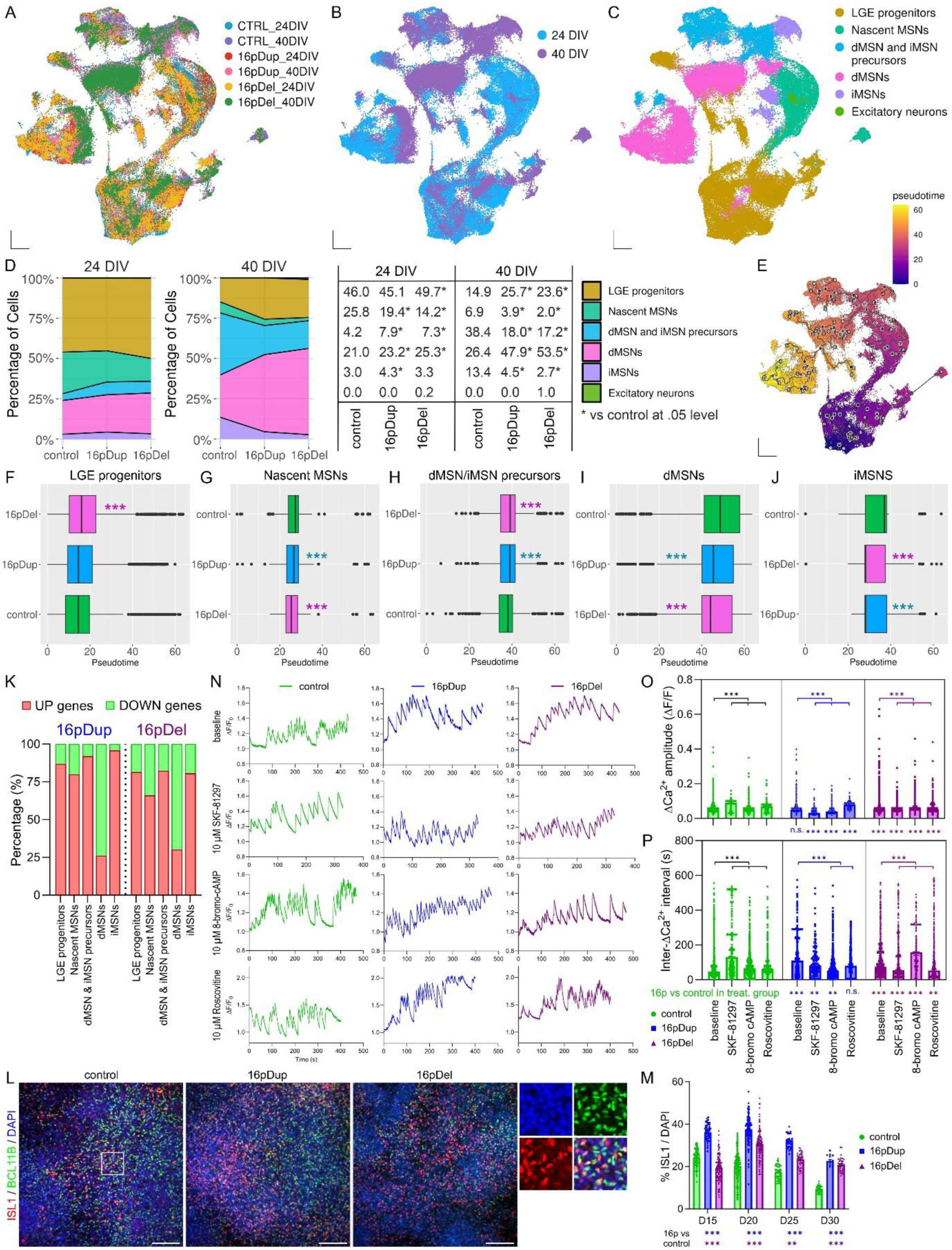
Bias toward dMSN fate in 16p11.2 cultures. (A–C) UMAP visualisations of control and 16p11.2dup/del MSN scRNA-seq data coloured by (A) sample, (B) time point and (C) cell type. (D) Cell population ratios at each time point depicted by stack plots (left) and compared by Pearson chi-square test followed by Bonferroni correction (right). (E) Trajectory analysis showing the pseudotime of striatal cells on UMAP. (F–J) Box plots showing the pseudotime of different cell types in iPSC-derived MSNs. (K) Quantification of upregulated and downregulated dMSN-specific DEGs in each cell type split by 16p11.2dup/del CNV. (L) Immunostaining for ISL1 and BCL11B and (M) quantification of ISL1^+^ cells in striatal cultures. (N) Representative traces of ΔCa^2+^ transients and quantification of (O) ΔCa^2+^ amplitude and (P) inter-ΔCa^2+^ intervals in control and CNV MSNs at baseline and with application of 10μM SKF-81927, 10μM 8-bromo-cAMP and 10μM Roscovitine. (M,O and P) Data represent the mean ± SEM with sample sizes summarised in **Table S4**. (F–J,M,O and P) Bonferroni corrected multiple comparison analysis with independent-samples median test (F–J,O and P) or Kruskal-Wallis test (M) were used for statistical analyses. **p < 0.01, ***p < 0.001. Scale bars: 100 μm in (L). See also **Figure S4** and **Tables S3 and S4**.

As dMSNs are typically born earlier than iMSNs^39,47^, this could stem from either delayed iMSN fate induction or precocious dMSN specification, which we assessed next using trajectory analysis (**Fig 4E–J**). Mirroring changes in differentiation kinetics, 16p11.2del samples exhibited delayed LGE fate acquisition compared to controls (**Fig 4F**). Importantly, we demonstrate earlier specification of nascent MSN and both dMSN/iMSN fates in both 16p11.2 genotypes compared to controls (**Fig 4G–J**). Differential gene expression analyses revealed selective upregulation of dMSN genes across multiple cell populations in both 16p11.2 genotypes, including progenitors, nascent MSNs, subtype precursors and iMSN clusters (**Fig 4K**). For instance, known dMSN marker *GAP43* showed earlier, more pronounced expression in both genotypes, while the novel direct-pathway candidate gene *NRN1* was markedly higher in 16p11.2del MSN subtype precursors (28.5% vs. 5.5% cells in controls) and dMSNs (39.0% vs. 31.2%) (**Fig S4I and Table S3**). These data suggest that the observed dMSN enrichment may occur, at least in part, due to precocious direct-pathway fate acquisition in 16p11.2 MSNs.

Immunolabelling for ISL1, an early dMSN marker^48,49^, further corroborated preferential dMSN commitment in both genotypes. This was demonstrated by consistently increased proportions of ISL1^+^ cells in 16p11.2dup cultures compared to controls from early time points (15DIV) and a preferential rise in ISL1^+^ vs DARPP32^+^ MSNs at 30DIV (2.4-fold for ISL1 vs. 1.5-fold for DARPP32) (**Fig 4L and 4M**). Despite overall reduction of MSNs, 16p11.2del samples also showed elevated numbers of ISL1^+^ cells from 20DIV compared to controls (**Fig 4L and 4M**).

We next sought to assess the effects of dMSN overproduction on network activity by performing calcium imaging in 16p11.2 CNV and control cultures at 50DIV. Neurons were treated with drugs known to modulate dMSN-mediated calcium signalling in the striatum, including dMSN agonist SKF-81297, cAMP analogue 8-bromo-cAMP, and CDK5 inhibitor roscovitine^50,51^. Examination of spontaneous activity-evoked Ca^2+^ transients (ΔCa^2+^) showed that both 16p11.2 genotypes exhibited reduced and less frequent ΔCa^2+^ at baseline compared to controls (**Fig 4N–4O**). Intriguingly, direct pathway-specific stimulation resulted in decreased inter-ΔCa^2+^ intervals in 16p11.2 MSNs in all treatment groups while controls displayed longer intervals, suggesting that dMSN transcriptional shifts correlate with CNV-dependent hyperexcitability or increased responsiveness of dMSNs (**Fig 4O**). Although effects on ΔCa^2+^ amplitude were more variable between genotypes, most treatments induced increases in the amplitude compared to baseline activity (**Fig 4P**). Collectively, these data underscore how reciprocal 16p11.2 CNVs can reprogram both fate and function in striatal neurons, potentially converging on similar basal ganglia circuit deficits in NDDs.

## Discussion

Here, we demonstrate high-fidelity transcriptional alignment of iPSC-derived dMSNs and iMSNs to human foetal and adult striatal signatures^39–41^, underscoring the suitability of our Activin A-patterned MSNs for probing how genetic risk variants disrupt striatal circuits in NDDs. Human iPSC-based models are particularly valuable for capturing early neuropathology in psychiatric conditions with a neurodevelopmental component^52,53^. By leveraging this framework, we establish a platform to investigate the pathophysiology of NDDs involving basal ganglia dysfunction—an area of growing interest given its role in diverse mental health conditions^2–7^.

Focusing on 16p11.2 CNVs, we uncover a dual phenomenon: (1) reciprocal shifts in cell-cycle kinetics that alter the timing and amplitude of MSN production, and (2) a shared bias favouring dMSN fate in both duplication and deletion genotypes. The divergent striatal neurogenesis deficits mirror phenotypes reported in CNV carriers and rodent models, where 16p11.2del and 16p11.2dup often manifest as macrocephaly versus microcephaly and hyper-versus hypo-proliferation^54–56^. Our data further demonstrate that these CNVs not only produce global neurogenesis phenotypes but also impose MSN-specific deficits likely contributing to striatal pathology and associated behavioural symptoms in carriers. The reciprocal cell-cycle changes reported here echo findings in cortical neurons derived from 16p11.2 iPSCs^20,57^. Moreover, similar cell-cycle and cortical neurogenesis disruptions have been documented in other NDD-associated CNVs or gene knock-out/overexpression models, such as 15q11.2, *CYFIP1*, *SETBP1* and *EZH1* among others^23–27^, indicating that multiple risk mutations converge on early-stage deficits in proliferation and neuronal maturation.

Notably, a key implication of our results is that dMSN enrichment may underlie specific motor and cognitive symptoms frequently noted in 16p11.2dup-associated schizophrenia and other conditions linked with both 16p11.2 genotypes, such as ASD, intellectual disability, and ADHD^5,8–13,58–61^. In our cultures, this dMSN overproduction correlates with enhanced firing frequency upon direct-pathway stimulation, indicating a direct link between transcriptional dMSN biases (e.g. *GAP43*, *ISL1, NRN1*) and functional shifts at the circuit level. Because dMSNs facilitate excitatory output in the basal ganglia^62^, an imbalance in dMSN/iMSN proportions could disrupt movement planning or reward-related behaviours. Moreover, 16p11.2-associated NDDs often present with features overlapping with ASD and schizophrenia^11,12,42^, disorders where basal ganglia involvement is increasingly recognised. Intriguingly, hyperexcitable striatal circuits have been implicated in repetitive behaviours in ASD models^59,63^, and abnormal dMSN/iMSN ratios can alter motivational states relevant to psychosis and schizophrenia^5,6^. Here, we provide direct evidence in human cells that 16p11.2 CNVs lead to hyperresponsive dMSNs, a novel discovery concordant with clinical observations of hyperactivity, repetitive behaviours and motor incoordination in 16p11.2 carriers and mouse models^14–17,64–66^. Despite evidence in human 16p11.2 neurons mainly arising from cortical, dopaminergic, or hippocampal lineages, multiple recent studies also report CNV-driven increases in neuronal excitability, inhibitory deficits causing global circuit hyperactivity, altered synaptic gene expression, and abnormal neurite outgrowth^19,22,57,67–69^. Our demonstration that striatal neurons exhibit analogous vulnerabilities broadens this narrative, indicating that 16p11.2 perturbations at different nodes across the motor and reward pathways may synergise to produce complex neurodevelopmental phenotypes.

Moving forward, dissecting which genes within 16p11.2 orchestrate proliferative or subtype-specific programmes in striatal neurons will be critical. Our transcriptomic analyses reveal that one gene set (*BCKDK*, *KIF22*, *PPP4C*, *QPRT*, *HIRIP3*) is upregulated in striatal progenitors, while another (*ASPHD1*, *KCTD13*, *SEZ6L2*, *TLCD3B*, *YPEL3*) is enriched in dMSN/iMSN precursors (**Table S2**). Some genes, such as *PPP4C* and *KIF22*, contribute to cell-cycle progression and mitotic spindle formation^70–72^, whereas *KCTD13* and *SEZ6L2* have been linked to synaptic function and dendritic development^73–77^. Our data indicate that manipulating these gene pathways could modulate proliferation rates, dMSN bias, or both. From a therapeutic standpoint, by showing that 16p11.2-dependent transcriptional changes bear functional relevance, we pave the way for targeted drug screening aimed at normalising dMSN excitability and subtype balance in NDDs where basal ganglia dysfunction is a central feature. Overall, our findings sharpen the genotype-to-phenotype links in 16p11.2 syndromes and bolster the potential of iPSC-derived MSNs as a robust platform for disease modelling, mechanism-focused research, and therapeutic development.

## Resource availability

### Lead contact

Further information and requests for resources and reagents should be directed to and will be fulfilled by the lead contact, Marija Fjodorova.

### Materials availability

All unique reagents generated in this study are available from the lead contact with a completed materials transfer agreement.

### Data and code availability

Sequencing data reported in this paper are available with the SRA accession number PRJNA1228584 and will be publicly available upon publication. This study does not report original algorithms. Access to additional information generated in this study can be obtained upon request from the lead contact.

## Acknowledgements

We would like to thank all the children and their family members who took part in this study through the Cardiff University Rare Variant Research Programme, as well as all the support we have had from NHS medical genetic clinics, and support charities, including Unique. We also thank Mrs Olena Petter for iPSC reprograming and the core laboratory team of the Division of Psychological Medicine and Clinical Neurosciences for DNA sample management and genotyping. We are grateful to the families at the participating Simons Searchlight sites as well as the Simons Searchlight Consortium, formerly the Simons VIP Consortium. We appreciate obtaining access to iPSC lines on SFARI Base used in this work. RNA sequencing analysis was performed using the computational facilities of the Advanced Research Computing@Cardiff (ARCCA) Division, Cardiff University. This research was funded by the Medical Research Council as part of the ‘Therapeutic target validation in mental health’ call (MR/S037667/1 to R.B., M.L., J.G., M.v.d.B.), via MR/R022429/1 to M.L. and a lectureship by the Hodge Foundation to M.F.. J.H. and M.v.d.B. receive additional support via MR/W028395/1 and R.B. was supported by the NextGeneration EU (NGEU), the Ministry of University and Research (MUR), National Recovery and Resilience Plan (NRRP), project MNESYS (PE0000006) – A multiscale integrated approach to the study of the nervous system in health and disease (DN. 1553 11 October 2022).

## Author contributions

M.F. and M.L. conceived the study and designed the experiments. M.F., P.P. and O.P. carried out the hiPSC experiments and M.F. analysed all data. M.F., P.P., N.N.V. and S.D. carried out the scRNA-seq experiments. M.F. and P.P. performed scRNA-seq data analysis with support from Z.L.. I.M., J.H., J.G., J.H., M.V.D.B. and R.B. provided 16p11.2 CNV carrier samples and contributed to general discussions throughout the work. M.F. and M.L. wrote the paper. All authors edited and approved the paper.

## Methods

### Experimental model and subject details

#### Cell lines

Fibroblasts from one adult female subject and peripheral blood mononuclear cells (PBMCs) from one male teenage subject carrying 16p11.2dup were reprogrammed into induced pluripotent stem cells (iPSCs) using the CytoTune-IPS 2.0 Sendai reprogramming kit (ThermoFisher). Same protocol was applied to reprogram PBMCs from three adult female subjects carrying 16p11.2del, as well as fibroblasts from two adult healthy subjects (one male and one female). Male hiPSCs HPSI0114i-kolf_2-C1 were commercially available from EBiSC and female hiPSCs SV0002291 carrying 16p11.2dup were obtained from Simons Foundation^19,78^. The iPSC lines used in this study were regularly tested for mycoplasma contamination.

Generation and genetic manipulation of human iPSC were approved by Cardiff University and HSE (GMO130/669). Ethical approvals were granted by the NHS Health Research Authority and Healthcare Research Wales, Manchester University NHS foundation trust research office (IRAS project ID 254310) and the East Midlands - Leicester Central Research Ethics Committee (REC reference 19/EM/0287). All experiments were performed in accordance with relevant guidelines and regulations, including the Declaration of Helsinki.

### Method Details

#### Stem cell culture and differentiation

All hPSCs were cultured on Matrigel coated plates in E8 media (ThermoFisher) in a humidified 37°C incubator with 5% CO_2_. Media was changed daily, and cells passaged mechanically with 0.02% EDTA at 80% confluence. MSNs were obtained and maintained as described previously^31,37^. In short, neural differentiation was initiated at 80% confluence (0DIV) by switching E8 medium to DMEM-F12/Neurobasal (2:1; Thermo Fisher Scientific) supplemented with N2 (Thermo Fisher Scientific) and vitamin A-free B27 (Thermo Fisher Scientific) [referred to thereafter as N2B27]. For the first 7DIV, cultures were supplemented with SB431542 (10µM, Tocris), LDN-193189 (100nM, StemGent) and dorsomorphin dihydrochloride (200nM, Tocris). Splitting was done en bloc using EDTA at 6DIV onto fibronectin-coated (Merck Millipore) plates at a ratio of 2:3. MSN fate was induced from 7DIV by supplementing N2B27 medium with activin A (25ng/ml, Cell Guidance Systems) until 30DIV. Cultures were split again at 16DIV onto poly-D-lysine hydrobromide (PDL, Merck Sigma-Aldrich) and laminin (Merck Sigma-Aldrich) co-coated plates at a density of 150k cells/cm2. B27 with vitamin A (Thermo Fisher Scientific) was used from 20DIV onwards. Differentiating MSNs were maintained in N2B27 supplemented with BDNF, GDNF (10ng/ml, PeproTech) from 20DIV to aid neuronal maturation and survival.

#### scRNA-seq and data processing

For scRNA-seq, differentiating MSNs were collected and dissociated into single cells at 15, 18, 24 and 40DIV. Briefly, cells were treated with Accutase (ThermoFisher) for 10–15 minutes at 37°C. The suspension was centrifuged at 200g for 5 minutes to remove supernatant. Cells were then re-suspended in PBS and passed through a 40μm filter. Three biological replicates of the same stage were pooled together. For quality control, the cell viability was confirmed to be above 80%. Cells were loaded onto the 10X Chromium Single Cell Platform and libraries were constructed using the Single Cell 3’ Library v3.1 kit (10X Genomics) as per manufacturer’s protocol. The libraries were sequenced on the Illumina Novaseq 6000 instrument aiming for a depth of approximately 150,000 reads/cell using paired-end sequencing.

To obtain count matrices, FASTQ files were mapped to the human genome GRCh38.84 (hg38, provided by 10X Genomics) following Cell Ranger (v6.1.1) workflow with default parameters. Downstream analysis was performed using the R package Seurat (v4.4.0)^79^. For quality control, cells with detected genes >300, UMI count <150,000, and the fraction of mitochondrial genes <10% were kept as high-quality cells. The normalisation and variance stabilisation of molecular count data were performed using sctransform (v2) function in Seurat^80^. This function was used to also remove confounding sources of variation such as mitochondrial mapping percentage.

After performing dimensional reduction, the batch effect was removed using harmony (v1.0.3) on the top 50 dimensions^81^. The cells were then visualised using Uniform Manifold Approximation and Projection (UMAP) embedding. The top 50 dimensions were used to identify neighbours of cells and clusters with a resolution of 0.4. Differential gene expression analyses were performed using the FindAllMarkers() function in Seurat with min.pct=0.25, logfc.threshold=0.25 parameters using wilcox test. Genes with adjusted *p* value<0.01 were defined as statistically significant. Cluster annotations were determined by analysing cluster-specific gene enrichment for cell type markers from previously annotated reference atlases of various primary human brain regions^39,40^. We performed trajectory analysis using the monocle3 package (v 1.2.9) with default parameters^82^, and the forebrain progenitor cluster was chosen as the root node to order cells. Cell cycle phase scores were calculated in the CellCycleScoring() function based on the expression of canonical G2/M and S phase markers and stored in object meta data, along with the predicted classification of each cell in either G2M, S or G1 phase.

To compare the transcriptome similarity with published human foetal and adult striatal scRNAseq data^39–41^, we performed anchor-based label transfer implemented in Seurat. The anchors between the query and reference datasets were identified and filtered using the FindTransferAnchors() function with the top 50 dimensions. Then, the MapQuery() wrapper function was used to project human brain data onto our data or vice versa to visualise cell type and developmental stage similarities between datasets. Same workflow was applied to assess similarities between Activin A- and SHH-patterned MSNs^43^.

To directly compare iPSC-derived MSNs to striatal neurons^39^, we integrated scRNA-seq datasets using anchor-based method in Seurat. After applying sctransform() function, the top 3000 variable features were used to determine anchors and perform integration. After performing dimensional reduction, the top 50 dimensions were used to identify neighbours of cells and clusters with a resolution of 0.5. Clusters were annotated as described above and cell type marker genes that were conserved across the two datasets were identified using the FindConservedMarkers() function in Seurat.

#### Immunocytochemistry

Cultured cells were rinsed in PBS and fixed in 3.7% PFA for 15 minutes at 4°C. For nuclear antigen detection an additional fixation with methanol gradient was performed, which includes 5 minutes each in 33% and 66% methanol at room temperature followed by 100% methanol for 20 minutes at −20°C. Cultures were then returned to PBS via inverse gradient. Cells were then permeabilized in 0.3% Triton-X-100 solution in PBS (PBS-T) and then blocked in PBS-T containing 1% BSA and 3% donkey serum. Cells were incubated with primary antibodies in blocking solution overnight at 4°C. Following three PBS-T washes, Alexa-Fluor secondary antibodies (Thermo Fisher Scientific) were added at 1:1000 in blocking solution for 1hr at ambient temperature in the dark. Cells were stained with DAPI at 1:1000 (Thermo Fisher Scientific). Images were taken on a PerkinElmer OperaPhenix microscope. Quantification was carried out in Cell Profiler (cellprofiler.org) or manually using ImageJ (imagej.net) by examining 10–15 randomly selected fields of view from 2–3 technical replicates performed in 3 independent differentiations (biological replicates) unless stated otherwise (**Table S4**). The following primary antibodies were used for the immunofluorescence studies: rabbit anti-FOXG1 (Abcam #ab18259, 1:1000), rabbit anti-GSX2 (Merck Millipore #ABN162, 1:500), rat anti-BCL11B (Abcam #ab18465, 1:500), rabbit anti-DARPP32 (Abcam #ab40802, 1:500), mouse anti-FOXP1 (Abcam #ab32010, 1:800), rabbit anti-FOXP2 (Abcam #ab16046, 1:500), mouse anti-NEUN (Sigma-Aldrich #MAB377, 1:500), rabbit anti-Ki67 (Abcam #ab15580, 1:500), mouse anti-ASCL1 (BD Biosciences #556604, 1:500), mouse anti-ISL1 (Developmental Studies Hybridoma Bank #39.4D5, 1:500).

#### EdU labelling and detection

Cells in S-phase were labelled using the Click-iT EdU Assay kit. For cumulative EdU labelling assay required for calculating cell cycle length, cells were incubated with EdU for increasing periods of time of 0, 1.5, 3, 6, 9, 12, 15, 18, 21, 24 and 30hr. Cells were then fixed in 3.7% PFA for 15 minutes at 4°C. EdU detection was carried out as per manufacturer’s protocol. Twenty fields each from 4 technical replicates and 2–4 independent differentiations were counted at each time point of the EdU cumulative labelling (**Table S4**). The percentages of EdU^+^ cells in each genotype were plotted against time. Cell cycle length was determined using the Nowakowski equation as depicted in **Figure 3L**^45^. A linear regression was performed, and the values of the slope (m) and y-intercept (b) of the linear equation were used to calculate cell cycle length (T_C_) and S-phase length (T_S_) as follows:

f(t) = (GF/ T_C_)*t + GF*(T_S_ /T_C_), where GF = growth fraction in cell population, T_C_ = GF/m and T_S_ = (b* T_C_)/GF.

#### Live-cell calcium imaging

Neuronal cultures were incubated with a Ca^2+^-sensitive probe Fluo-4-AM (Thermo Fisher Scientific) at a final concentration of 5µM for 30 minutes at 37°C. The solution was then replaced with pre-warmed artificial cerebrospinal fluid (aCSF) solution [2mM CaCl_2_, 142mM NaCl, 2.5mM KCl, 1mM MgCl_2_, 10mM HEPES, 30mM D-glucose – all from Merck Sigma-Aldrich]. Where indicated aCSF was supplemented with one of the following: 10µM SKF-81297, 10µM 8-bromo-cAMP, 10µM Roscovitine (all from Tocris). All drug solutions were prepared fresh on the day, pre-warmed to 37°C and delivered/aspirated to/from cultured cells via a pump delivery system. Spontaneous activity- and drug-evoked Ca^2+^ transients were recorded on a Zeiss AG Axio Observer D1 inverted microscope. Subsequent data processing was performed using FluoroSNNAP algorithm in MATLAB (MathWorks) according to the protocol^83^. Individual traces were measured from 100–200 neurons per each of the 5–7 fields of view from 2–3 technical replicates and 2 independent experiments (**Table S4**).

#### Statistical analysis

RNA sequencing data was analysed separately as described above. SPSS Statistics 29 software (IBM) was used for all other statistical analyses. All quantified data were plotted in Prism 10 (GraphPad Software) and are reported as mean ± SEM with sample sizes for each test summarised in **Table S4**. Normal distribution of data was assessed using the Shapiro-Wilk test and homogeneity of variance analysed with Levene’s test as part of testing assumptions for parametric tests. Statistical analyses were performed using either one- or two-way ANOVA tests followed by Tukey HSD correction for multiple comparisons if applicable. Alternatively, non-parametric Kruskal-Wallis test or independent-samples median test followed by Bonferroni correction was used for data that did not meet the above assumptions. Nominal data were analysed using Pearson chi-square test followed by Bonferroni correction for multiple comparisons. Results were considered statistically significant at p<0.05. Randomisation was used to assign experimental conditions and data collection was always done in parallel with controls.

## References

1. Reiner, A., Albin, R.L., Anderson, K.D., Damato, C.J., Penney, J.B., and Young, A.B. (1988). Differential loss of striatal projection neurons in Huntington disease. Proceedings of the National Academy of Sciences of the United States of America 85, 5733–5737. 10.1073/pnas.85.15.5733.

2. Gamazon, E.R., Zwinderman, A.H., Cox, N.J., Denys, D., and Derks, E.M. (2019). Multi-tissue transcriptome analyses identify genetic mechanisms underlying neuropsychiatric traits. Nature Genetics 51, 933–940. 10.1038/s41588-019-0409-8.

3. Huckins, L.M., Dobbyn, A., Ruderfer, D.M., Hoffman, G., Wang, W., Pardinas, A.F., Rajagopal, V.M., Als, T.D., Nguyen, H.T., Girdhar, K., et al. (2019). Gene expression imputation across multiple brain regions provides insights into schizophrenia risk. Nature Genetics 51, 659–674. 10.1038/s41588-019-0364-4.

4. Fuccillo, M. (2016). Striatal Circuits as a Common Node for Autism Pathophysiology. FRONTIERS IN NEUROSCIENCE 10, 27. 10.3389/fnins.2016.00027.

5. Vicente, A.M., Martins, G.J., and Costa, R.M. (2020). Cortico-basal ganglia circuits underlying dysfunctional control of motor behaviors in neuropsychiatric disorders. Current Opinion in Genetics & Development 65, 151–159. 10.1016/j.gde.2020.05.042.

6. Skene, N.G., Bryois, J., Bakken, T.E., Breen, G., Crowley, J.J., Gaspar, H.A., Giusti-Rodriguez, P., Hodge, R.D., Miller, J.A., Muñoz-Manchado, A.B., et al. (2018). Genetic identification of brain cell types underlying schizophrenia. Nature Genetics 50, 825–833. 10.1038/s41588-018-0129-5.

7. Duncan, L., Li, T., Salem, M., Li, W., Mortazavi, L., Senturk, H., Shahverdizadeh, N., Vesuna, S., Shen, H., Yoon, J., et al. (2025). Mapping the cellular etiology of schizophrenia and complex brain phenotypes. NATURE NEUROSCIENCE. 10.1038/s41593-024-01834-w.

8. Robinson, J.E., and Gradinaru, V. (2018). Dopaminergic dysfunction in neurodevelopmental disorders: recent advances and synergistic technologies to aid basic research. Current Opinion in Neurobiology 48, 17–29. 10.1016/j.conb.2017.08.003.

9. Di, Y., Diao, Z., Zheng, Q., Li, J., Cheng, Q., Li, Z., Fang, S., Wang, H., Wei, C., Zheng, Q., et al. (2022). Differential Alterations in Striatal Direct and Indirect Pathways Mediate Two Autism-like Behaviors in Valproate-Exposed Mice. JOURNAL OF NEUROSCIENCE 42, 7833–7847. 10.1523/JNEUROSCI.0623-22.2022.

10. Niarchou, M., Chawner, S., Doherty, J.L., Maillard, A.M., Jacquemont, S., Chung, W.K., Green-Snyder, L., Bernier, R.A., Goin-Kochel, R.P., Hanson, E., et al. (2019). Psychiatric disorders in children with 16p11.2 deletion and duplication. Translational Psychiatry 9, 8, 8. 10.1038/s41398-018-0339-8.

11. Weiss, L.A., Shen, Y.P., Korn, J.M., Arking, D.E., Miller, D.T., Fossdal, R., Saemundsen, E., Stefansson, H., Ferreira, M.A.R., Green, T., et al. (2008). Association between microdeletion and microduplication at 16p11.2 and autism. New England Journal of Medicine 358, 667–675. 10.1056/NEJMoa075974.

12. Leone, R., Zuglian, C., Brambilla, R., and Morella, I. (2024). Understanding copy number variations through their genes: a molecular view on 16p11.2 deletion and duplication syndromes. FRONTIERS IN PHARMACOLOGY 15, 1407865. 10.3389/fphar.2024.1407865.

13. Gur, R., Bearden, C., Jacquemont, S., Swillen, A., van Amelsvoort, T., van den Bree, M., Vorstman, J., Sebat, J., Ruparel, K., Gallagher, R., et al. (2025). Neurocognitive profiles of 22q11.2 and 16p11.2 deletions and duplications. MOLECULAR PSYCHIATRY 30, 379–387. 10.1038/s41380-024-02661-y.

14. Portmann, T., Yang, M., Mao, R., Panagiotakos, G., Ellegood, J., Dolen, G., Bader, P.L., Grueter, B.A., Goold, C., Fisher, E., et al. (2014). Behavioral Abnormalities and Circuit Defects in the Basal Ganglia of a Mouse Model of 16p11.2 Deletion Syndrome. Cell Reports 7, 1077–1092. 10.1016/j.celrep.2014.03.036.

15. Arbogast, T., Ouagazzal, A.M., Chevalier, C., Kopanitsa, M., Afinowi, N., Migliavacca, E., Cowling, B.S., Birling, M.C., Champy, M.F., Reymond, A., and Herault, Y. (2016). Reciprocal Effects on Neurocognitive and Metabolic Phenotypes in Mouse Models of 16p11.2 Deletion and Duplication Syndromes. Plos Genetics 12, e1005709. 10.1371/journal.pgen.1005709.

16. Rein, B., and Yan, Z. (2020). 16P11.2 Copy Number Variations and Neurodevelopmental Disorders. TRENDS IN NEUROSCIENCES 43, 886–901. 10.1016/j.tins.2020.09.001.

17. Cunningham, A., Hall, J., Owen, M., and van den Bree, M. (2021). Coordination difficulties, IQ and psychopathology in children with high-risk copy number variants. PSYCHOLOGICAL MEDICINE 51, 290–299, PII S0033291719003210. 10.1017/S0033291719003210.

18. Grissom, N.M., McKee, S.E., Schoch, H., Bowman, N., Havekes, R., O’Brien, W.T., Mahrt, E., Siegel, S., Commons, K., Portfors, C., et al. (2018). Male-specific deficits in natural reward learning in a mouse model of neurodevelopmental disorders. Molecular Psychiatry 23, 544–555. 10.1038/mp.2017.184.

19. Deshpande, A., Yadav, S., Dao, D., Wu, Z., Hokanson, K., Cahill, M., Wiita, A., Jan, Y., Ullian, E., and Weiss, L. (2017). Cellular Phenotypes in Human iPSC-Derived Neurons from a Genetic Model of Autism Spectrum Disorder. CELL REPORTS 21, 2678–2687. 10.1016/j.celrep.2017.11.037.

20. Connacher, R., Williams, M., Prem, S., Yeung, P.L., Matteson, P., Mehta, M., Markov, A., Peng, C., Zhou, X.F., McDermott, C.R., et al. (2022). Autism NPCs from both idiopathic and CNV 16p11.2 deletion patients exhibit dysregulation of proliferation and mitogenic responses. Stem Cell Reports 17, 1380–1394. 10.1016/j.stemcr.2022.04.019.

21. Roth, J., Muench, K., Asokan, A., Mallett, V., Gai, H., Verma, Y., Weber, S., Charlton, C., Fowler, J., Loh, K., et al. (2020). 16p11.2 microdeletion imparts transcriptional alterations in human iPSC-derived models of early neural development. ELIFE 9, e58178. 10.7554/eLife.58178.

22. Fetit, R., Barbato, M., Theil, T., Pratt, T., and Price, D. (2023). 16p11.2 deletion accelerates subpallial maturation and increases variability in human iPSC-derived ventral telencephalic organoids. DEVELOPMENT 150, dev201227. 10.1242/dev.201227.

23. Gracia-Diaz, C., Zhou, Y., Yang, Q., Maroofian, R., Espana-Bonilla, P., Lee, C., Zhang, S., Padilla, N., Fueyo, R., Waxman, E., et al. (2023). Gain and loss of function variants in EZH1 disrupt neurogenesis and cause dominant and recessive neurodevelopmental disorders. NATURE COMMUNICATIONS 14. 10.1038/s41467-023-39645-5.

24. Cardo, L., de la Fuente, D., and Li, M. (2023). Impaired neurogenesis and neural progenitor fate choice in a human stem cell model of SETBP1 disorder. MOLECULAR AUTISM 14, 8. 10.1186/s13229-023-00540-x.

25. De La Fuente, D., Tamburini, C., Stonelake, E., Andrews, R., Hall, J., Owen, M., Linden, D., Pocklington, A., and Li, M. (2024). Impaired oxysterol-liver X receptor signaling underlies aberrant cortical neurogenesis in a stem cell model of neurodevelopmental disorder. Cell Reports 43, 113946.

26. Paulsen, B., Velasco, S., Kedaigle, A., Pigoni, M., Quadrato, G., Deo, A., Adiconis, X., Uzquiano, A., Sartore, R., Yang, S., et al. (2022). Autism genes converge on asynchronous development of shared neuron classes. NATURE 602, 268-+. 10.1038/s41586-021-04358-6.

27. Durak, O., Gao, F., Kaeser-Woo, Y., Rueda, R., Martorell, A., Nott, A., Liu, C., Watson, L., and Tsai, L. (2016). Chd8 mediates cortical neurogenesis via transcriptional regulation of cell cycle and Wnt signaling. NATURE NEUROSCIENCE 19, 1477–1488. 10.1038/nn.4400.

28. Mariani, J., Coppola, G., Zhang, P., Abyzov, A., Provini, L., Tomasini, L., Amenduni, M., Szekely, A., Palejev, D., Wilson, M., et al. (2015). FOXG1-Dependent Dysregulation of GABA/Glutamate Neuron Differentiation in Autism Spectrum Disorders. CELL 162, 375–390. 10.1016/j.cell.2015.06.034.

29. Marchetto, M., Belinson, H., Tian, Y., Freitas, B., Fu, C., Vadodaria, K., Beltrao-Braga, P., Trujillo, C., Mendes, A., Padmanabhan, K., et al. (2017). Altered proliferation and networks in neural cells derived from idiopathic autistic individuals. MOLECULAR PSYCHIATRY 22, 820–835. 10.1038/mp.2016.95.

30. Cambray, S., Arber, C., Little, G., Dougalis, A.G., de Paola, V., Ungless, M.A., Li, M., and Rodriguez, T.A. (2012). Activin induces cortical interneuron identity and differentiation in embryonic stem cell-derived telencephalic neural precursors. Nature Communications 3, 841. 10.1038/ncomms1817.

31. Arber, C., Precious, S.V., Cambray, S., Risner-Janiczek, J.R., Kelly, C., Noakes, Z., Fjodorova, M., Heuer, A., Ungless, M.A., Rodriguez, T.A., et al. (2015). Activin A directs striatal projection neuron differentiation of human pluripotent stem cells. Development 142, 1375–1386. 10.1242/dev.117093.

32. Fjodorova, M., Noakes, Z., and Li, M. (2020). A role for TGFβ signalling in medium spiny neuron differentiation of human pluripotent stem cells. Neuronal Signaling 4, NS20200004.

33. Nicoleau, C., Varela, C., Bonnefond, C., Maury, Y., Bugi, A., Aubry, L., Viegas, P., Bourgois-Rocha, F., Peschanski, M., and Perrier, A.L. (2013). Embryonic Stem Cells Neural Differentiation Qualifies the Role of Wnt/beta-Catenin Signals in Human Telencephalic Specification and Regionalization. Stem Cells 31, 1763–1774. 10.1002/stem.1462.

34. Aubry, L., Bugi, A., Lefort, N., Rousseau, F., Peschanski, M., and Perrier, A.L. (2008). Striatal progenitors derived from human ES cells mature into DARPP32 neurons in vitro and in quinolinic acid-lesioned rats. Proceedings of the National Academy of Sciences of the United States of America 105, 16707–16712. 10.1073/pnas.0808488105.

35. Ma, L., Hu, B., Liu, Y., Vermilyea, S.C., Liu, H., Gao, L., Sun, Y., Zhang, X., and Zhang, S.-C. (2012). Human Embryonic Stem Cell-Derived GABA Neurons Correct Locomotion Deficits in Quinolinic Acid-Lesioned Mice. Cell Stem Cell 10, 455–464. 10.1016/j.stem.2012.01.021.

36. Carri, A.D., Onorati, M., Lelos, M.J., Castiglioni, V., Faedo, A., Menon, R., Camnasio, S., Vuono, R., Spaiardi, P., Talpo, F., et al. (2013). Developmentally coordinated extrinsic signals drive human pluripotent stem cell differentiation toward authentic DARPP-32(+) medium-sized spiny neurons. Development 140, 301–312. 10.1242/dev.084608.

37. Fjodorova, M., Louessard, M., Li, Z., De La Fuente, D.C., Dyke, E., Brooks, S.P., Perrier, A.L., and Li, M. (2019). CTIP2-Regulated Reduction in PKA-Dependent DARPP32 Phosphorylation in Human Medium Spiny Neurons: Implications for Huntington Disease. Stem Cell Reports 13, 448–457. 10.1016/j.stemcr.2019.07.015.

38. Fjodorova, M., Noakes, Z., de la Fuente, D., Errington, A., and Li, M. (2023). Dysfunction of cAMP-Protein Kinase A-Calcium Signaling Axis in Striatal Medium Spiny Neurons: A Role in Schizophrenia and Huntington’s Disease Neuropathology. BIOLOGICAL PSYCHIATRY: GLOBAL OPEN SCIENCE 3, 418–429. 10.1016/j.bpsgos.2022.03.010.

39. Bocchi, V.D., Conforti, P., Vezzoli, E., Besusso, D., Cappadona, C., Lischetti, T., Galimberti, M., Ranzani, V., Bonnal, R.J.P., De Simone, M., et al. (2021). The coding and long noncoding single-cell atlas of the developing human fetal striatum. Science 372, 591-+, eabf5759. 10.1126/science.abf5759.

40. Shi, Y.C., Wang, M.D., Mi, D., Lu, T., Wang, B.S., Dong, H., Zhong, S.J., Chen, Y.Q., Sun, L., Zhou, X., et al. (2021). Mouse and human share conserved transcriptional programs for interneuron development. Science 374, 1342-+, eabj6641. 10.1126/science.abj6641.

41. Lee, H., Fenster, R.J., Pineda, S.S., Gibbs, W.S., Mohammadi, S., Davila-Velderrain, J., Garcia, F.J., Therrien, M., Novis, H.S., Gao, F., et al. (2020). Cell Type-Specific Transcriptomics Reveals that Mutant Huntingtin Leads to Mitochondrial RNA Release and Neuronal Innate Immune Activation. Neuron 107, 891-+. 10.1016/j.neuron.2020.06.021.

42. Niarchou, M., Chawner, S., Doherty, J.L., Maillard, A.M., Jacquemont, S., Chung, W.K., Green-Snyder, L., Bernier, R.A., Goin-Kochel, R.P., Hanson, E., et al. (2019). Psychiatric disorders in children with 16p11.2 deletion and duplication. Translational Psychiatry 9, 8. 10.1038/s41398-018-0339-8.

43. Conforti, P., Bocchi, V.D., Campus, I., Scaramuzza, L., Galimberti, M., Lischetti, T., Talpo, F., Pedrazzoli, M., Murgia, A., Ferrari, I., et al. (2022) *In vitro*-derived medium spiny neurons recapitulate human striatal development and complexity at single-cell resolution. Cell Reports Methods 2. 10.1016/j.crmeth.2022.100367.

44. Zhang, Z.Z., Wei, S., Du, H., Su, Z.H., Wen, Y., Shang, Z.C., Song, X.L., Xu, Z.J., You, Y., and Yang, Z.G. (2019). *Zfhx3* is required for the differentiation of late born D1-type medium spiny neurons. Experimental Neurology 322, 113055. 10.1016/j.expneurol.2019.113055.

45. Nowakowski, R., Lewin, S., and Miller, M. (1989). BROMODEOXYURIDINE IMMUNOHISTOCHEMICAL DETERMINATION OF THE LENGTHS OF THE CELL-CYCLE AND THE DNA-SYNTHETIC PHASE FOR AN ANATOMICALLY DEFINED POPULATION. JOURNAL OF NEUROCYTOLOGY 18, 311–318.

46. Arai, Y., Pulvers, J.N., Haffner, C., Schilling, B., Nüsslein, I., Calegari, F., and Huttner, W.B. (2011). Neural stem and progenitor cells shorten S-phase on commitment to neuron production. Nature Communications 2, 154. 10.1038/ncomms1155.

47. Shang, Z.C., Yang, L., Wang, Z.W., Tian, Y., Gao, Y.J., Su, Z.H., Guo, R.L., Li, W.W., Liu, G.P., Li, X.S., et al. (2022). The transcription factor *Zfp503* promotes the D1 MSN identity and represses the D2 MSN identity. Frontiers in Cell and Developmental Biology 10, 948331. 10.3389/fcell.2022.948331.

48. Ehrman, L.A., Mu, X.Q., Waclaw, R.R., Yoshida, Y., Vorhees, C.V., Klein, W.H., and Campbell, K. (2013). The LIM homeobox gene *Isl1* is required for the correct development of the striatonigral pathway in the mouse. Proceedings of the National Academy of Sciences of the United States of America 110, E4026–E4035. 10.1073/pnas.1308275110.

49. Lu, K.M., Evans, S.M., Hirano, S., and Liu, F.C. (2014). Dual role for *Islet-1* in promoting striatonigral and repressing striatopallidal genetic programs to specify striatonigral cell identity. Proceedings of the National Academy of Sciences of the United States of America 111, E168–E177. 10.1073/pnas.1319138111.

50. Tang, T.-S., and Bezprozvanny, I. (2004). Dopamine receptor-mediated Ca2+ signaling in striatal medium spiny neurons. Journal of Biological Chemistry 279, 42082–42094. 10.1074/jbc.M407389200.

51. Bibb, J.A., Snyder, G.L., Nishi, A., Yan, Z., Meijer, L., Fienberg, A.A., Tsai, L.H., Kwon, Y.T., Girault, J.A., Czernik, A.J., et al. (1999). Phosphorylation of DARPP-32 by Cdk5 modulates dopamine signalling in neurons. Nature 402, 669–671. 10.1038/45251.

52. Hannon, E., Spiers, H., Viana, J., Pidsley, R., Burrage, J., Murphy, T.M., Troakes, C., Turecki, G., O’Donovan, M.C., Schalkwyk, L.C., et al. (2016). Methylation QTLs in the developing brain and their enrichment in schizophrenia risk loci. Nature Neuroscience 19, 48-+. 10.1038/nn.4182.

53. Gulsuner, S., Walsh, T., Watts, A.C., Lee, M.K., Thornton, A.M., Casadei, S., Rippey, C., Shahin, H., Nimgaonkar, V.L., Go, R.C.P., et al. (2013). Spatial and Temporal Mapping of De Novo Mutations in Schizophrenia to a Fetal Prefrontal Cortical Network. Cell 154, 518–529. 10.1016/j.cell.2013.06.049.

54. Qureshi, A.Y., Mueller, S., Snyder, A.Z., Mukherjee, P., Berman, J.I., Roberts, T.P.L., Nagarajan, S.S., Spiro, J.E., Chung, W.K., Sherr, E.H., et al. (2014). Opposing Brain Differences in 16p11.2 Deletion and Duplication Carriers. Journal of Neuroscience 34, 11199–11211. 10.1523/jneurosci.1366-14.2014.

55. McCarthy, S.E., Makarov, V., Kirov, G., Addington, A.M., McClellan, J., Yoon, S., Perkins, D.O., Dickel, D.E., Kusenda, M., Krastoshevsky, O., et al. (2009). Microduplications of 16p11.2 are associated with schizophrenia. Nature Genetics 41, 1223–U1285. 10.1038/ng.474.

56. Pucilowska, J., Vithayathil, J., Tavares, E., Kelly, C., Karlo, J., and Landreth, G. (2015). The 16p11.2 Deletion Mouse Model of Autism Exhibits Altered Cortical Progenitor Proliferation and Brain Cytoarchitecture Linked to the ERK MAPK Pathway. JOURNAL OF NEUROSCIENCE 35, 3190–3200. 10.1523/JNEUROSCI.4864-13.2015.

57. Urresti, J., Zhang, P., Moran-Losada, P., Yu, N., Negraes, P., Trujillo, C., Antaki, D., Amar, M., Chau, K., Pramod, A., et al. (2021). Cortical organoids model early brain development disrupted by 16p11.2 copy number variants in autism. MOLECULAR PSYCHIATRY 26, 7581–7581. 10.1038/s41380-021-01289-6.

58. Abbott, A., Linke, A., Nair, A., Jahedi, A., Alba, L., Keown, C., Fishman, I., and Müller, R. (2018). Repetitive behaviors in autism are linked to imbalance of corticostriatal connectivity: a functional connectivity MRI study. SOCIAL COGNITIVE AND AFFECTIVE NEUROSCIENCE 13, 32–42. 10.1093/scan/nsx129.

59. Gandhi, T., and Lee, C. (2021). Neural Mechanisms Underlying Repetitive Behaviors in Rodent Models of Autism Spectrum Disorders. FRONTIERS IN CELLULAR NEUROSCIENCE 14, 592710. 10.3389/fncel.2020.592710.

60. Ferhat, A., Verpy, E., Biton, A., Forget, B., De Chaumont, F., Mueller, F., Le Sourd, A., Coqueran, S., Schmitt, J., Rochefort, C., et al. (2023). Excessive self-grooming, gene dysregulation and imbalance between the striosome and matrix compartments in the striatum of Shank3 mutant mice. FRONTIERS IN MOLECULAR NEUROSCIENCE 16, 1139118. 10.3389/fnmol.2023.1139118.

61. Schuetze, M., Park, M.T.M., Cho, I.Y.K., MacMaster, F.P., Chakravarty, M.M., and Bray, S.L. (2016). Morphological Alterations in the Thalamus, Striatum, and Pallidum in Autism Spectrum Disorder. Neuropsychopharmacology 41, 2627–2637. 10.1038/npp.2016.64.

62. Lanciego, J., Luquin, N., and Obeso, J. (2012). Functional Neuroanatomy of the Basal Ganglia. COLD SPRING HARBOR PERSPECTIVES IN MEDICINE 2, a009621. 10.1101/cshperspect.a009621.

63. Peixoto, R.T., Chantranupong, L., Hakim, R., Levasseur, J., Wang, W.G., Merchant, T., Gorman, K., Budnik, B., and Sabatini, B.L. (2019). Abnormal Striatal Development Underlies the Early Onset of Behavioral Deficits in Shank3B(-/-) Mice. Cell Reports 29, 2016–2027. 10.1016/j.celrep.2019.10.021.

64. Rein, B., Tan, T., Yang, F., Wang, W., Williams, J., Zhang, F., Mills, A., and Yan, Z. (2021). Reversal of synaptic and behavioral deficits in a 16p11.2 duplication mouse model via restoration of the GABA synapse regulator Npas4. MOLECULAR PSYCHIATRY 26, 1967–1979. 10.1038/s41380-020-0693-9.

65. Wang, Q., Qin, J., Chen, Y., Yu, Y., Xing, Y., Wang, Y., Yang, L., Lu, S., Geng, L., Shi, W., et al. (2023). 16p11.2 CNV gene Doc2a functions in neurodevelopment and social behaviors through interaction with Secretagogin. CELL REPORTS 42, 112691. 10.1016/j.celrep.2023.112691.

66. Kim, J., Vanrobaeys, Y., Kelvington, B., Peterson, Z., Baldwin, E., Gaine, M., Nickl-Jockschat, T., and Abel, T. (2024). Dissecting 16p11.2 hemi-deletion to study sex-specific striatal phenotypes of neurodevelopmental disorders. MOLECULAR PSYCHIATRY 29, 1310–1321. 10.1038/s41380-024-02411-0.

67. Forrest, M., Dos Santos, M., Piguel, N., Wang, Y., Hawkins, N., Bagchi, V., Dionisio, L., Yoon, S., Simkin, D., Martin-de-Saavedra, M., et al. (2023). Rescue of neuropsychiatric phenotypes in a mouse model of 16p11.2 duplication syndrome by genetic correction of an epilepsy network hub. NATURE COMMUNICATIONS 14, 825. 10.1038/s41467-023-36087-x.

68. Parnell, E., Culotta, L., Forrest, M., Jalloul, H., Eckman, B., Loizzo, D., Horan, K., Dos Santos, M., Piguel, N., Tai, D., et al. (2023). Excitatory Dysfunction Drives Network and Calcium Handling Deficits in 16p11.2 Duplication Schizophrenia Induced Pluripotent Stem Cell-Derived Neurons. BIOLOGICAL PSYCHIATRY 94, 153–163. 10.1016/j.biopsych.2022.11.005.

69. Sundberg, M., Pinson, H., Smith, R., Winden, K., Venugopal, P., Tai, D., Gusella, J., Talkowski, M., Walsh, C., Tegmark, M., and Sahin, M. (2021). 16p11.2 deletion is associated with hyperactivation of human iPSC-derived dopaminergic neuron networks and is rescued by RHOA inhibition in vitro. NATURE COMMUNICATIONS 12, 2897. 10.1038/s41467-021-23113-z.

70. Blaker-Lee, A., Gupta, S., McCammon, J., De Rienzo, G., and Sive, H. (2012). Zebrafish homologs of genes within 16p11.2, a genomic region associated with brain disorders, are active during brain development, and include two deletion dosage sensor genes. DISEASE MODELS & MECHANISMS 5, 834–851. 10.1242/dmm.009944.

71. Kawaue, H., Matsubara, T., Nagano, K., Ikedo, A., Rojasawasthien, T., Yoshimura, A., Nakatomi, C., Imai, Y., Kakuta, Y., Addison, W., and Kokabu, S. (2024). KIF22 regulates mitosis and proliferation of chondrocyte cells. ISCIENCE 27, 110151. 10.1016/j.isci.2024.110151.

72. Liao, F., Hsiao, W., Lin, Y., Chan, Y., and Huang, C. (2016). T cell proliferation and adaptive immune responses are critically regulated by protein phosphatase 4. CELL CYCLE 15, 1073–1083. 10.1080/15384101.2016.1156267.

73. Yaguchi, H., Yabe, I., Takahashi, H., Watanabe, M., Nomura, T., Kano, T., Matsumoto, M., Nakayama, K., Watanabe, M., and Hatakeyama, S. (2017). Sez6l2 regulates phosphorylation of ADD and neuritogenesis. BIOCHEMICAL AND BIOPHYSICAL RESEARCH COMMUNICATIONS 494, 234–241. 10.1016/j.bbrc.2017.10.047.

74. Escamilla, C., Filonova, I., Walker, A., Xuan, Z., Holehonnur, R., Espinosa, F., Liu, S., Thyme, S., López-García, I., Mendoza, D., et al. (2017). Kctd13 deletion reduces synaptic transmission via increased RhoA. NATURE 551, 227-+. 10.1038/nature24470.

75. Qiu, W., Luo, S., Ma, S., Saminathan, P., Li, H., Gunnersen, J., Gelbard, H., and Hammond, J. (2021). The Sez6 Family Inhibits Complement by Facilitating Factor I Cleavage of C3b and Accelerating the Decay of C3 Convertases. FRONTIERS IN IMMUNOLOGY 12, 607641. 10.3389/fimmu.2021.607641.

76. Boonen, M., Staudt, C., Gilis, F., Oorschot, V., Klumperman, J., and Jadot, M. (2016). Cathepsin D and its newly identified transport receptor SEZ6L2 can modulate neurite outgrowth. JOURNAL OF CELL SCIENCE 129, 557–568. 10.1242/jcs.179374.

77. Lorenzo, S., Nalesso, V., Chevalier, C., Birling, M., and Herault, Y. (2021). Targeting the RHOA pathway improves learning and memory in adult Kctd13 and 16p11.2 deletion mouse models. MOLECULAR AUTISM 12, 1. 10.1186/s13229-020-00405-7.

78. Hildebrandt, M., Reuter, M., Wei, W., Tayebi, N., Liu, J., Sharmin, S., Mulder, J., Lesperance, L., Brauer, P., Mok, R., et al. (2019). Precision Health Resource of Control iPSC Lines for Versatile Multilineage Differentiation. STEM CELL REPORTS 13, 1126–1141. 10.1016/j.stemcr.2019.11.003.

79. Hao, Y., Hao, S., Andersen-Nissen, E., Mauck, W., Zheng, S., Butler, A., Lee, M., Wilk, A., Darby, C., Zager, M., et al. (2021). Integrated analysis of multimodal single-cell data. CELL 184, 3573-+. 10.1016/j.cell.2021.04.048.

80. Choudhary, S., and Satija, R. (2022). Comparison and evaluation of statistical error models for scRNA-seq. GENOME BIOLOGY 23, 27. 10.1186/s13059-021-02584-9.

81. Korsunsky, I., Millard, N., Fan, J., Slowikowski, K., Zhang, F., Wei, K., Baglaenko, Y., Brenner, M., Loh, P., and Raychaudhuri, S. (2019). Fast, sensitive and accurate integration of single-cell data with Harmony. NATURE METHODS 16, 1289-+. 10.1038/s41592-019-0619-0.

82. Cao, J., Spielmann, M., Qiu, X., Huang, X., Ibrahim, D., Hill, A., Zhang, F., Mundlos, S., Christiansen, L., Steemers, F., et al. (2019). The single-cell transcriptional landscape of mammalian organogenesis. NATURE 566, 496-+. 10.1038/s41586-019-0969-x.

83. Patel, T.P., Man, K., Firestein, B.L., and Meaney, D.F. (2015). Automated quantification of neuronal networks and single-cell calcium dynamics using calcium imaging. Journal of Neuroscience Methods 243, 26–38. 10.1016/j.jneumeth.2015.01.020.

