## Supplemental Information for "Direct pathway bias and altered striatal neurogenesis in human iPSC models of 16p11.2 CNVs: Evidence from single-cell and functional analyses"

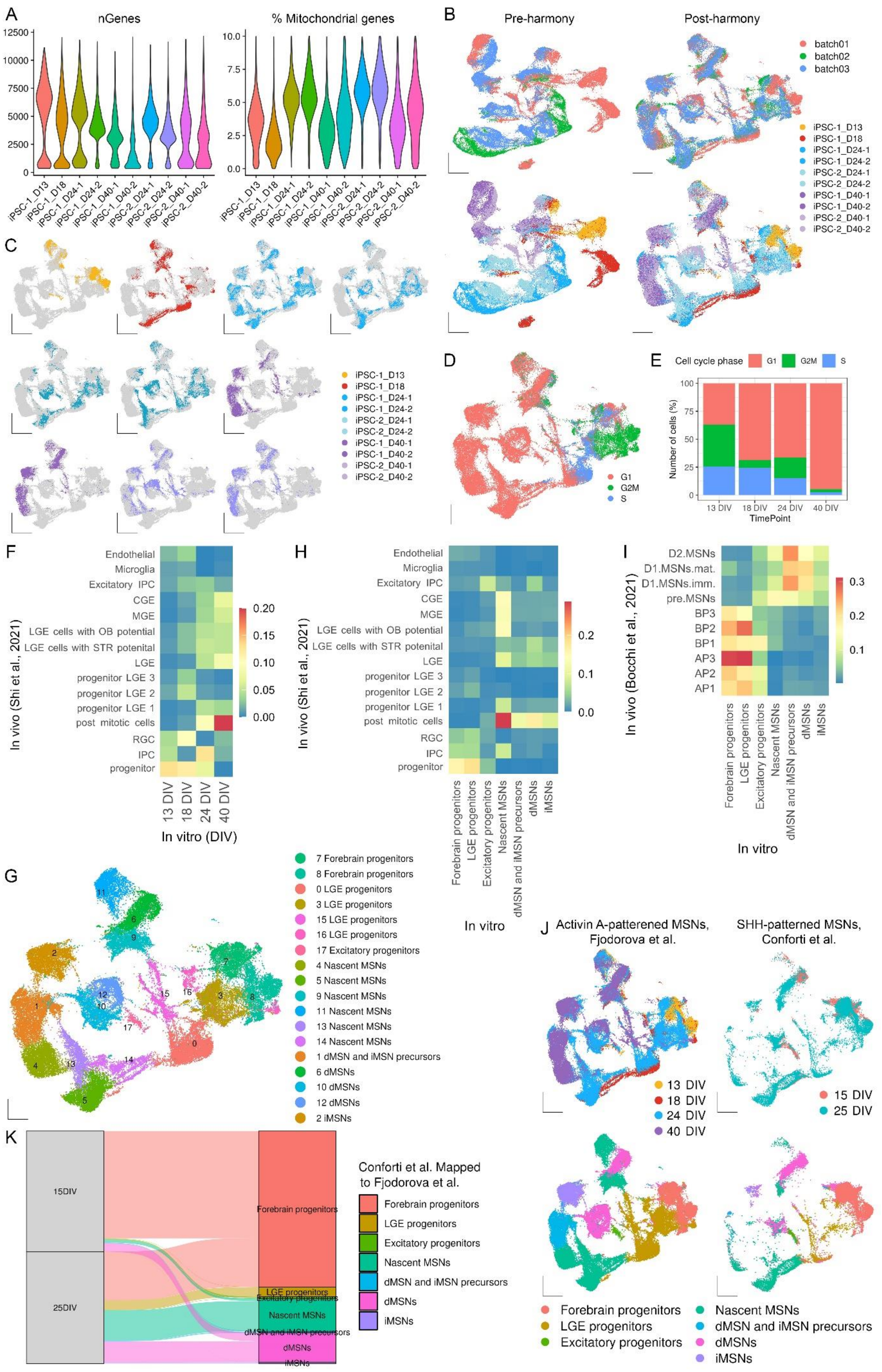
A
nGenes
% Mitochondrial genes
iPSC-1_D13
iPSC-1_D18
iPSC-1_D24-1
iPSC-1_D24-2
iPSC-1_D40-1
iPSC-1_D40-2
iPSC-2_D24-1
iPSC-2_D24-2
iPSC-2_D40-1
iPSC-2_D40-2
B
Pre-harmony
Post-harmony
batch01
batch02
batch03
C
D
G1
G2M
S
E
Cell cycle phase
Number of cells (%)
13 DIV
18 DIV
24 DIV
40 DIV
TimePoint
F
In vivo (Shi et al., 2021)
Endothelial
Microglia
Excitatory IPC
CGE
MGE
LGE cells with OB potential
LGE cells with STR potential
LGE
progenitor LGE 3
progenitor LGE 2
progenitor LGE 1
post mitotic cells
RGC
IPC
progenitor
In vitro (DIV)
H
Forebrain progenitors
LGE progenitors
Excitatory progenitors
Nascent MSNs
dMSN and iMSN precursors
dMSNs
iMSNs
In vitro
I
In vivo (Bocchi et al., 2021)
D2.MSNs
D1.MSNs.mat.
D1.MSNs.imm.
pre.MSNs
BP3
BP2
BP1
AP3
AP2
AP1
G
7 Forebrain progenitors
8 Forebrain progenitors
0 LGE progenitors
3 LGE progenitors
15 LGE progenitors
16 LGE progenitors
17 Excitatory progenitors
4 Nascent MSNs
5 Nascent MSNs
9 Nascent MSNs
11 Nascent MSNs
13 Nascent MSNs
14 Nascent MSNs
1 dMSN and iMSN precursors
6 dMSNs
10 dMSNs
12 dMSNs
2 iMSNs
J
Activin A-patterened MSNs, Fjodorova et al.
SHH-patterned MSNs, Conforti et al.
13 DIV
18 DIV
24 DIV
40 DIV
15 DIV
25 DIV
Forebrain progenitors
LGE progenitors
Excitatory progenitors
Nascent MSNs
dMSN and iMSN precursors
dMSNs
iMSNs
K
15DIV
25DIV
Conforti et al. Mapped to Fjodorova et al.
Forebrain progenitors
LGE progenitors
Excitatory progenitors
Nascent MSNs
dMSN and iMSN precursors
dMSNs
iMSNs

**Figure S1 Transcriptional profiling of Activin A-patterned human MSNs, related to Figure 1**

(A) Violin plots showing the number of genes (left) and percentage of mitochondrial genes (right) expressed in each sample. (B) UMAP plots showing the scRNA-seq data before (left) and after (right) batch effect correction with harmony coloured by batch (top) and sample (bottom). (C) UMAP embedding of sample profiles split by sample. (D) UMAP plot of single cells colour coded by predicted cell cycle phase. (E) The proportion of cells in each cell cycle phase at different stages of MSN differentiation. (F) Heatmap showing the similarity between different cell types in human foetal ganglionic eminence (9–18 PCW)[40] and different time points in MSN data. (G) UMAP embedding of scRNA-seq data in each Seurat cluster. (H–I) Heatmaps of transcriptome similarity between different cell types in Activin A-patterned MSNs and human foetal (H) ganglionic eminence (9–18 PCW)[40] or (I) lateral ganglionic eminence (7–11 PCW)[39]. (J) UMAP embedding of MapQuery projections of SHH-patterned MSNs (right)[43] onto scRNA-seq data colour coded by time point (top) and cell type (bottom). (K) Alluvial plot showing alignment of SHH-treated cultures at different stages of differentiation with different cell types in Activin A-patterned MSNs.

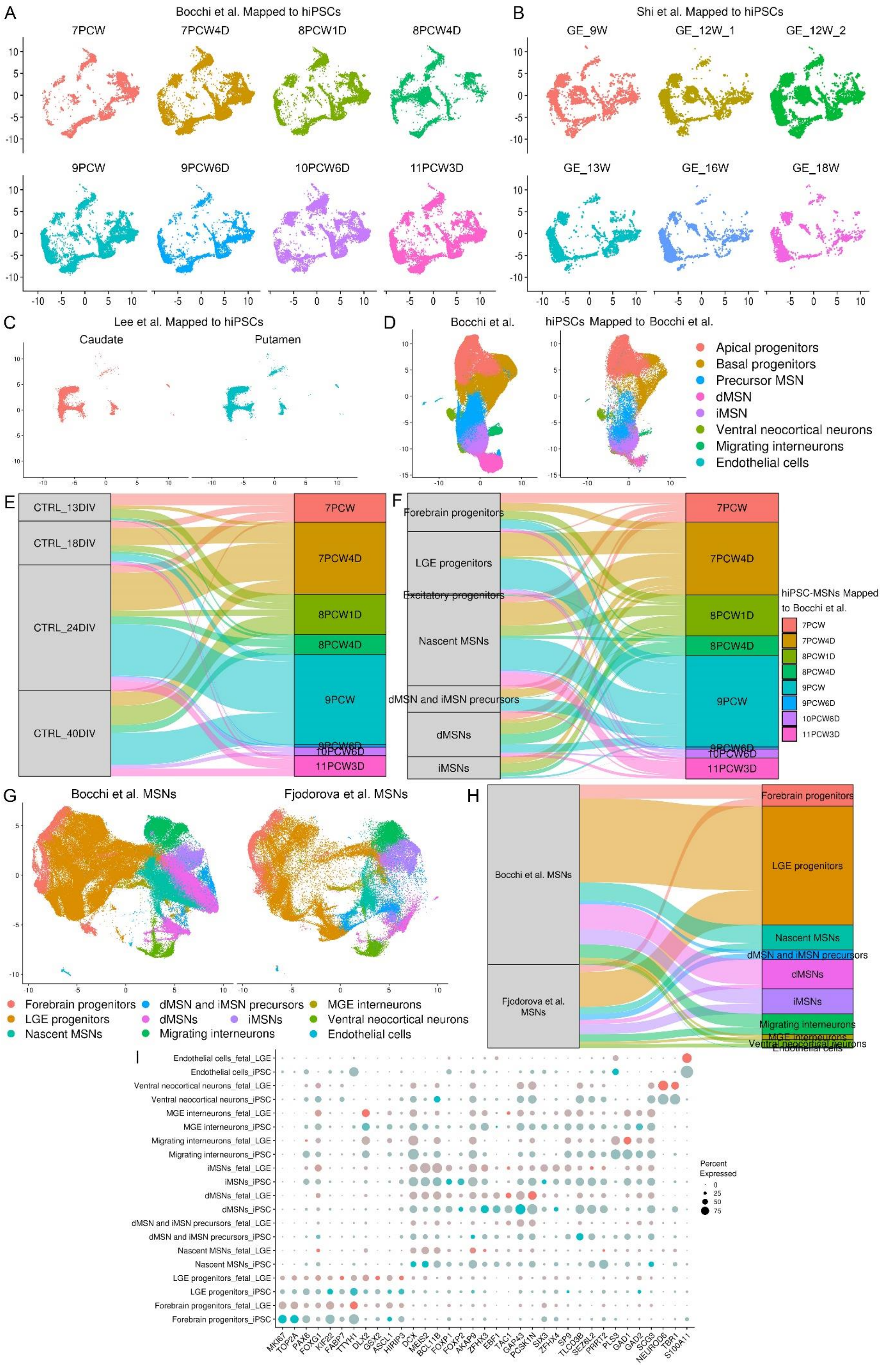
A
Bocchi et al. Mapped to hiPSCs
7PCW
7PCW4D
8PCW1D
8PCW4D
9PCW
9PCW6D
10PCW6D
11PCW3D
B
Shi et al. Mapped to hiPSCs
GE_9W
GE_12W_1
GE_12W_2
GE_13W
GE_16W
GE_18W
C
Lee et al. Mapped to hiPSCs
Caudate
Putamen
D
Bocchi et al.
hiPSCs Mapped to Bocchi et al.
Apical progenitors
Basal progenitors
Precursor MSN
dMSN
iMSN
Ventral neocortical neurons
Migrating interneurons
Endothelial cells
E
CTRL_13DIV
CTRL_18DIV
CTRL_24DIV
CTRL_40DIV
F
Forebrain progenitors
LGE progenitors
Excitatory progenitors
Nascent MSNs
dMSN and iMSN precursors
dMSNs
iMSNs
hiPSC-MSNs Mapped to Bocchi et al.
G
Bocchi et al. MSNs
Fjodorova et al. MSNs
Forebrain progenitors
LGE progenitors
Nascent MSNs
dMSN and iMSN precursors
dMSNs
iMSNs
Migrating interneurons
MGE interneurons
Ventral neocortical neurons
Endothelial cells
H
Bocchi et al. MSNs
Fjodorova et al. MSNs
I
Endothelial cells_fetal_LGE
Endothelial cells_iPSC
Ventral neocortical neurons_fetal_LGE
Ventral neocortical neurons_iPSC
MGE interneurons_fetal_LGE
MGE interneurons_iPSC
Migrating interneurons_fetal_LGE
Migrating interneurons_iPSC
iMSNs_fetal_LGE
iMSNs_iPSC
dMSNs_fetal_LGE
dMSNs_iPSC
dMSN and iMSN precursors_fetal_LGE
dMSN and iMSN precursors_iPSC
Nascent MSNs_fetal_LGE
Nascent MSNs_iPSC
LGE progenitors_fetal_LGE
LGE progenitors_iPSC
Forebrain progenitors_fetal_LGE
Forebrain progenitors_iPSC
MKI67 TOP2A PAX6 FOXG1 KIF22 FABP7 TTYH1 DLX2 GSX2 ASCL1 HIRIP3 DCX MEIS2 BCL11B FOXP1 FOXP2 AKAP9 ZFHX3 EBF1 TAC1 GAP43 PCSK1N SIX3 ZFHX4 SP9 TLCD3B SEZ6L2 PRRT2 PLS3 GAD1 GAD2 SCG3 NEUROD6 TBR1 S100A11
Percent Expressed
0
25
50
75

**Figure S2 Activin A protocol shows significant transcriptional similarity with human striatal neurons, related to Figure 2**

(A–C) UMAP embedding of MapQuery projections of human striatal cells at (A) 7–11 PCW[39], (B) 9–18 PCW[40] or (C) 32–88 years[41] onto scRNA-seq data split by sample. (D) UMAP embedding of MapQuery projections of iPSC-derived MSNs onto foetal striatal cells[39] coloured by cell type. (E–F) Alluvial plots showing alignment of MSNs by (E) differentiation stage or (F) cell type with foetal time points. (G) Integrated UMAP embedding of foetal striatal cells[39] and iPSC-derived MSNs coloured by cell type. (H) Alluvial plot showing the alignment of integrated datasets with cell types. (I) Dot plot showing the expression of signature conserved markers in each cell type of integrated data.

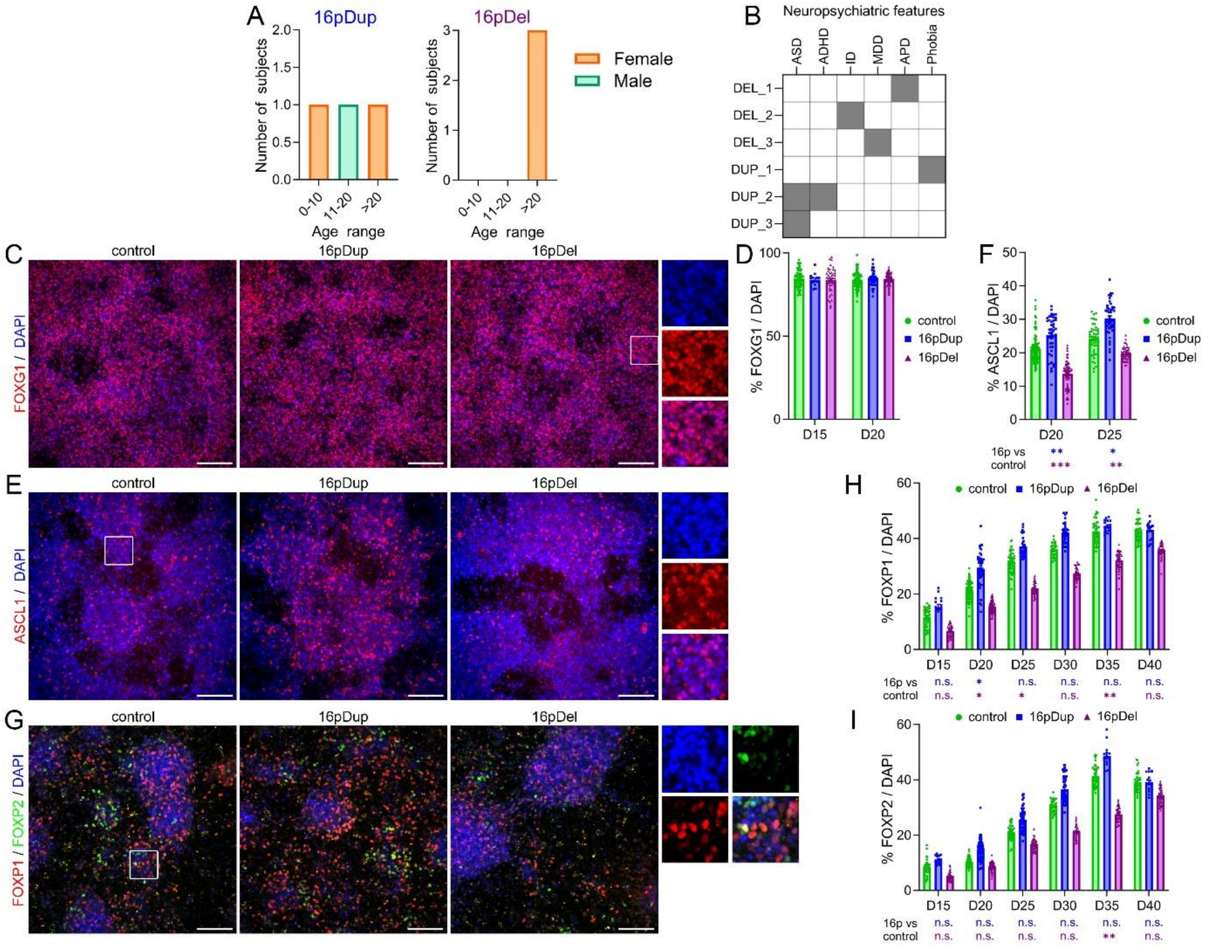

**Figure S3 Characterisation of control and 16p11.2dup/del iPSC-derived MSNs, related to Figure 3**

Summary of (A) subject demographics and (B) clinical features in 16p11.2 CNV carriers. (C) Immunostaining for FOXG1 and (D) quantification of marker$^+$ cells in differentiating control and 16p11.2dup/del iPSCs. (E) Immunostaining for ASCL1 and (F) quantification of ASCL1$^+$ cells in striatal neuron cultures. (G) Immunostaining for FOXP1 and FOXP2 in cultured MSNs. Quantification of (H) FOXP1$^+$ and (I) FOXP2$^+$ neurons in differentiating control and CNV cultures. (D, F, H and I) Data represent the mean ± SEM and sample sizes are summarised in **Table S4**. (D) Two-way ANOVA and (F, H and I) Bonferroni corrected multiple comparison analysis with Kruskal-Wallis test were used for statistical analyses. *p < 0.05, **p < 0.01, ***p < 0.001.

Scale bars: 100 μm in (C, E and G).

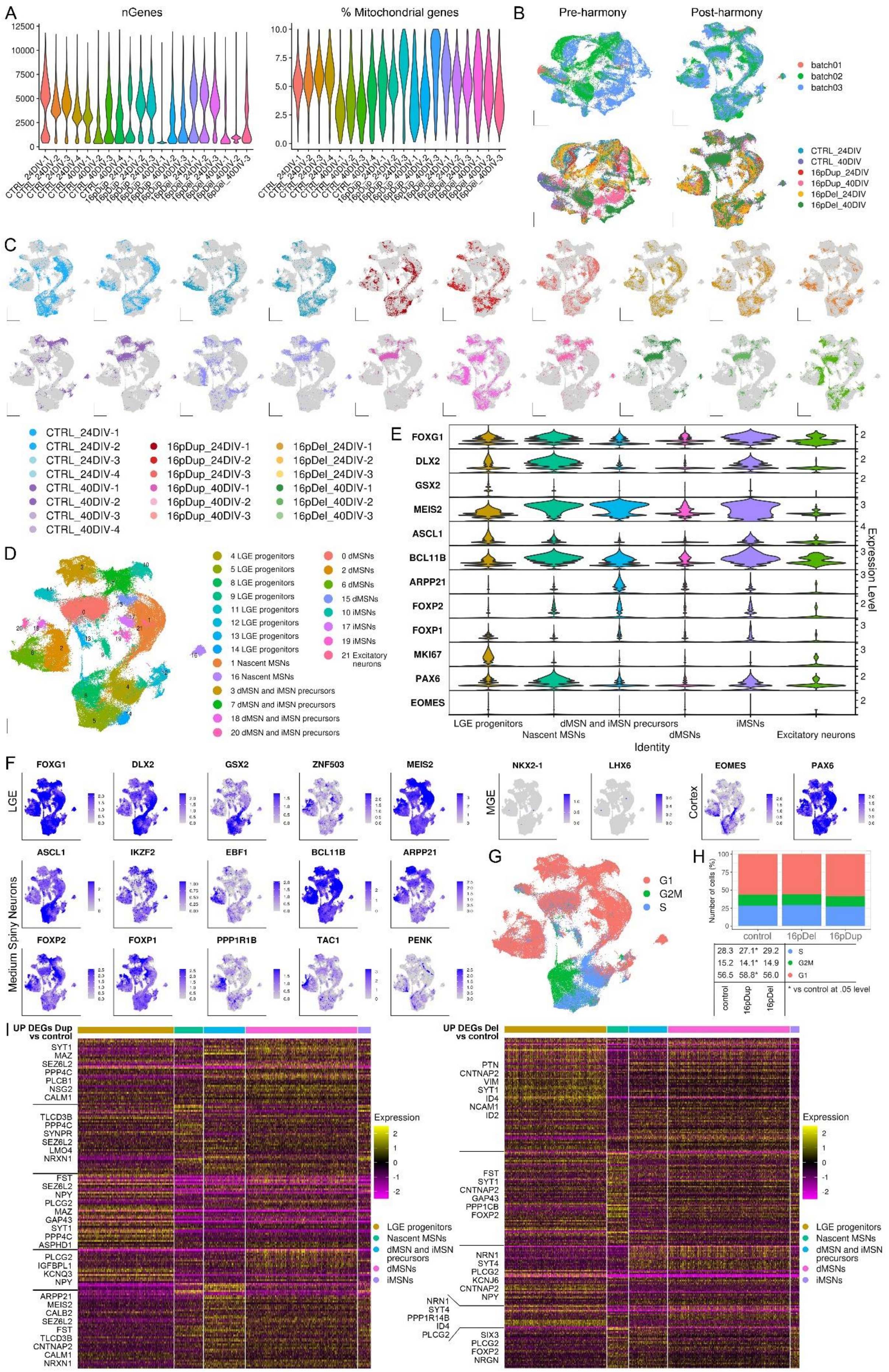
A
nGenes
% Mitochondrial genes
B
Pre-harmony
Post-harmony
batch01
batch02
batch03
CTRL_24DIV
CTRL_40DIV
16pDup_24DIV
16pDup_40DIV
16pDel_24DIV
16pDel_40DIV
C
CTRL_24DIV-1
CTRL_24DIV-2
CTRL_24DIV-3
CTRL_24DIV-4
CTRL_40DIV-1
CTRL_40DIV-2
CTRL_40DIV-3
CTRL_40DIV-4
16pDup_24DIV-1
16pDup_24DIV-2
16pDup_24DIV-3
16pDup_40DIV-1
16pDup_40DIV-2
16pDup_40DIV-3
16pDel_24DIV-1
16pDel_24DIV-2
16pDel_24DIV-3
16pDel_40DIV-1
16pDel_40DIV-2
16pDel_40DIV-3
D
4 LGE progenitors
5 LGE progenitors
8 LGE progenitors
9 LGE progenitors
11 LGE progenitors
12 LGE progenitors
13 LGE progenitors
14 LGE progenitors
1 Nascent MSNs
16 Nascent MSNs
3 dMSN and iMSN precursors
7 dMSN and iMSN precursors
18 dMSN and iMSN precursors
20 dMSN and iMSN precursors
0 dMSNs
2 dMSNs
6 dMSNs
15 dMSNs
10 iMSNs
17 iMSNs
19 iMSNs
21 Excitatory neurons
E
FOXG1
DLX2
GSX2
MEIS2
ASCL1
BCL11B
ARPP21
FOXP2
FOXP1
MKI67
PAX6
EOMES
Expression Level
LGE progenitors
Nascent MSNs
dMSN and iMSN precursors
dMSNs
iMSNs
Excitatory neurons
Identity
F
LGE
FOXG1
DLX2
GSX2
ZNF503
MEIS2
MGE
NKX2-1
LHX6
Cortex
EOMES
PAX6
Medium Spiny Neurons
ASCL1
IKZF2
EBF1
BCL11B
ARPP21
FOXP2
FOXP1
PPP1R1B
TAC1
PENK
G
G1
G2M
S
H
Number of cells (%)
control
16pDel
16pDup
28.3 27.1* 29.2 S
15.2 14.1* 14.9 G2M
56.5 58.8* 56.0 G1
control 16pDup 16pDel
* vs control at .05 level
I
UP DEGs Dup vs control
SYT1
MAZ
SEZ6L2
PPP4C
PLCB1
NSG2
CALM1
TLCD3B
PPP4C
SYNPR
SEZ6L2
LMO4
NRXN1
FST
SEZ6L2
NPY
PLCG2
MAZ
GAP43
SYT1
PPP4C
ASPHD1
PLCG2
IGFBPL1
KCNQ3
NPY
ARPP21
MEIS2
CALB2
SEZ6L2
FST
TLCD3B
CNTNAP2
CALM1
NRXN1
UP DEGs Del vs control
PTN
CNTNAP2
VIM
SYT1
ID4
NCAM1
ID2
FST
SYT1
CNTNAP2
GAP43
PPP1CB
FOXP2
NRN1
SYT4
PLCG2
KCNJ6
CNTNAP2
NPY
NRN1
SYT4
PPP1R14B
ID4
PLCG2
SIX3
PLCG2
FOXP2
NRGN
Expression
LGE progenitors
Nascent MSNs
dMSN and iMSN precursors
dMSNs
iMSNs

**Figure S4 Single-cell transcriptomic analysis of MSNs carrying 16p11.2 CNVs, related to Figure 4**

(A) Violin plots showing the number of genes (left) and percentage of mitochondrial genes (right) expressed in each sample. (B) UMAP plots showing the scRNA-seq data before (left) and after (right) batch effect correction with harmony coloured by batch (top) and sample (bottom). (C) UMAP embedding of sample profiles split by sample. (D) UMAP embedding of scRNA-seq data in each Seurat cluster. (E) Violin plot showing the expression of cell-type-specific genes in each cluster group. (F) Feature plots showing the expression levels of brain-region-specific genes in scRNA-seq data. (G) UMAP plot of single cells colour coded by predicted cell cycle phase. (H) The proportions of cells in each cell cycle phase for each genotype at 24DIV depicted by bar plot (top) and compared by Pearson chi-square test followed by Bonferroni correction (bottom), *$p < 0.05$. Sample sizes are summarised in **Table S4**. (I) Heatmaps showing DEGs upregulated in 16p11.2dup (left) and 16p11.2del (right) versus controls in each MSN cluster group.

### Supplemental table legends

**Table S1 (Related to Figure 1 and S1) – Summary of differential gene expression analysis and cluster ID assignment in control MSN dataset.** The first sheet provides an overview of assigned cluster IDs. The following sheet contains gene lists for individual seurat clusters (p_adj<0.01).

**Table S2 (Related to Figure 2 and S2) – Significantly upregulated genes by striatal cell population in control MSN dataset.**

**Table S3 (Related to Figure 4 and S4) – Summary of differential gene expression analyses in 16p11.2 CNV MSN dataset.** The first sheet contains upregulated gene lists for each cluster group (p_adj<0.01). The following two sheets contain significantly upregulated genes in each 16p11.2 genotype versus control by cluster group (p_adj<0.01).

**Table S4 (Related to Figures 3, 4, S3 and S4) – Additional information on sample numbers and cell lines.**
